# A statistical guide to the design of deep mutational scanning experiments

**DOI:** 10.1101/048892

**Authors:** Sebastian Matuszewski, Marcel E. Hildebrandt, Ana-Hermina Ghenu, Jeffrey D. Jensen, Claudia Bank

## Abstract

The characterization of the distribution of mutational effects is a key goal in evolutionary biology. Recently developed deep-sequencing approaches allow for accurate and simultaneous estimation of the fitness effects of hundreds of engineered mutations by monitoring their relative abundance across time points in a single bulk competition. Naturally, the achievable resolution of the estimated fitness effects depends on the specific experimental setup, the organism and type of mutations studied, and the sequencing technology utilized, among other factors. By means of analytical approximations and simulations, we provide guidelines for optimizing time-sampled deep-sequencing bulk competition experiments, focusing on the number of mutants, the sequencing depth, and the number of sampled time points. Our analytical results show that sampling more time points together with extending the duration of the experiment improves the achievable precision disproportionately as compared with increasing the sequencing depth, or reducing the number of competing mutants. Even if the duration of the experiment is fixed, sampling more time points and clustering these at the beginning and the end of the experiment increases experimental power, and allows for efficient and precise assessment of the entire range of selection coefficients. Finally, we provide a formula for calculating the 95%-confidence interval for the measurement error estimate, which we implement as an interactive web tool. This allows for quantification of the maximum expected *a priori* precision of the experimental setup, as well as for a statistical threshold for determining deviations from neutrality for specific selection coefficient estimates.

## Introduction

**M**utations provide the fuel for evolutionary change, and their fitness effects critically influence the course and dynamics of evolution. The distribution of fitness effects (DFE) lies at the heart of many evolutionary concepts, such as the genetic basis of complex traits (Eyre-Walker 2010) and diseases (Keightley and Eyre-Walker 2010), the rate of adaptation to a new environment (Gerrish and Lenski 1998; Orr 1998, Orr 2005b), the maintenance of genetic variation (Charlesworth *et al.* 1995), and the relative importance of selection and drift in molecular evolution (Ohta 1977,Ohta 1992; Kimura 1979). Unsurprisingly, considerable effort has been devoted, both empirically (e.g., Sawyer et al. 2003; Sousa *et al.* 2012; Gordo and Campos 2013; Bernet and Elena 2015) and theoretically (e.g., Gillespie 1983; Orr 2005a; Martin and Lenormand 2006b; Rice *et al.* 2015; Connallon and Clark 2015) to assess the fraction of all possible mutations that are beneficial, neutral, or deleterious. Until recently, the two main approaches (Eyre-Walker and Keightley 2007; Hietpas *et al.* 2011) for assessing the distribution of fitness effects (DFE) have been based either on the analysis of polymorphism and divergence data (Jensen *et al.* 2008; Keightley and Eyre-Walker 2010; Schneider *et al.* 2011) or on laboratory evolution studies in which spontaneously occurring mutations are followed for many generations (Imhof and Schlotterer 2001; Rozen *et al.* 2002; Halligan and Keightley 2010; Frenkel *et al.* 2014). However, the complex action and interaction of evolutionary forces within and between individuals and the environment makes accurate estimation of fitness effects of single mutations difficult (Orr 2009).

Recently, an alternative option to study mutational effects on a large scale has emerged from the field of biophysics: deep mutational scanning (DMS; Fowler *et al.* 2010; Hietpas *et al.* 2011; Fowler and Fields 2014). This approach is typically focused on a specific region of the genome for which a large library of mutants is created, either through random or systematic mutagenesis. The effects of the mutants are subsequently assessed by sequencing, with the readout yielding the relative frequencies of each mutant through time (obtained either directly, or via sequence tags). This results in a high-precision snapshot of local mutational effects without the influence of genome-wide interactions (e.g., epistasis) and environmental fluctuations.

DMS provides various advantages over traditional approaches of deriving DFEs from polymorphism and laboratory-evolution data. Firstly, it is not confounded by sampling bias (i.e., lethal mutations can also be observed) because the entire spectrum of pre-engineered or random mutations is introduced into a controlled and identical genetic background rather than waiting for mutations to appear and survive stochastic loss (Rokyta *et al.* 2005; Orr 2009). Secondly, the short time frame of the experiment and the large library size minimize the influence of secondary mutations, which eliminates the challenges imposed by epistasis and linked selection. Finally, bulk competition ensures that all mutants experience the same environment.

A DMS approach termed *EMPIRIC* (Hietpas *et al.* 2011) has been most prevalently studied with respect to estimation of the DFE and its application to evolutionary questions. *EMPIRIC* allows simultaneous estimation of the fitness of systematically engineered mutations in a given protein region. Mutants are constructed by transformation of pre-constructed plasmid mutant libraries, each representing one of all total point mutations from the focal protein region; these then undergo bulk competition for a number of generations. Fitness is determined by assessing relative growth rates from the relative abundance of each mutant, which is obtained from deep sequence data for a number of time points.

To date, *EMPIRIC* has been applied to yeast (*Saccharomyces cerevisiae*) to illuminate the DFE of all point mutations in Ubiq-uitin (Roscoe *et al.* 2013) and Hsp90 (Hietpas *et al.* 2011) across different environments, to quantify the amount and strength of epistatic interactions within a region of Hsp90 (Bank *et al.* 2014), and to assess a large intragenic fitness landscape in Hsp90. Recently, this approach has been extended to human influenza A virus to study the DFE in a region of the Neuraminidase protein containing a known drug-resistant locus. This opens the door for studying the mechanistic features underlying drug resistance and for determining potential future resistance mutations in viral populations (Jiang *et al.* 2015).

It has been demonstrated that the *EMPIRIC* approach is highly reproducible across replicate experiments and shows strong correspondence with selection coefficient estimates from binary competitions (Hietpas *et al.* 2011, 2013), resulting in precise estimates of selection coefficients (Bank *et al.* 2014). However, the attainable precision strongly depends on the experimental setup, in particular on the number of mutants considered, the number of time samples taken, and the sequencing depth. Furthermore all these factors need to be determined before the experiment and are constrained by the scientific question at hand, and additional limitations imposed by time and budget. The aim of this paper is to provide a statistical framework for *a priori* optimization of the experimental setup for future DMS studies (for an alternative approach see Kowalsky *et al.* (2015)).

Our model has been originally inspired by the *EMPIRIC*approach, but our predictions can be readily applied to any experiment that meets the following requirements (see Table 1 for further examples):

1. All studied mutants are present at large copy number at the beginning of the experiment (such that all mutants will be sampled sufficiently at later time points; usually on the order of 10^2^).
2. The population size is always kept lower than the carrying capacity (e.g., through serial dilution, or in a chemostat), such that mutants grow approximately exponentially (i.e., log-linearly) throughout the experiment.
3. Population size and sample size (for sequencing, or in case of serial passaging) are large compared with the number of mutants and sequencing depth.
4. Populations are sampled by deep sequencing (or fluorescence counting) at two or more time points, and individual mutant frequencies are assessed either directly or via sequence tags.

Thus, the statistical guidelines derived in the following can in principle be directly applied to experiments using new genome editing approaches based on CRISPR/Cas9 (Jinek *et al.* 2012), ZFN (Chen *et al.* 2011) and TALEN (Joung and Sander 2013) which constitute particularly exciting and promising new means for assessing the selective effects of new mutations (i.e., the DFE), but equally pertain to traditional binary competition experiments to assess relative growth rates. Note however, that DMS studies in which the functional capacity of a (mutant) protein (i.e., the protein fitness) cannot be directly related to organismal fitness (for a recent review on the topic see Boucher *et al.* 2016) do not adhere to the statistical framework presented here. Examples include the following recent DMS studies, which were based on fluorescence (as in Sarkisyan *et al.* 2016), antibiotic resistance (e.g., Jacquier *et al.* 2013; Firnberg *et al.* 2014), and binding selection using protein display technologies (Fowler *et al.* 2010; Whitehead *et al.* 2012; Olson *et al.* 2014).

Here, we derive analytical approximations for the variance and the mean squared error (MSE) of the estimators for the selection coefficients obtained by (log-)linear regression. We describe how measurement error decreases with the number of sampling time points and the number of sequencing reads, and how increasing the number of mutants generally increases the MSE. Based on these results, we derive the length of the 95%-confidence interval as an *a priori* measure of maximum attainable precision under a given experimental setup. Furthermore, we demonstrate that sampling more time points together with extending the duration of the experiment improves the achievable precision disproportionately as compared with increasing the sequencing depth. However, even if the duration of the experiment is fixed, sampling more time points and clustering these at the beginning and the end of the experiment increases experimental power and allows for efficient and precise assessment of selection coefficients of strongly deleterious as well as nearly neutral mutants. When applying our statistical framework to a data set of 568 engineered mutations from Hsp90 in *Saccharomyces cerevisiae*, we find that the experimental error is well predicted as long as the experimental requirements (see above) are met. To ease application of our results to future experiments, we provide an interactive online calculator (available onhttps://evoldynamics.org/tools).

**Table 1.** List of published DMS studies assessing mutant growth rates in accordance with our statistical model; arranged by number of time points sampled.

| Reference | Number of time points | Number of mutants | Total number of sequence reads (millions) | Reproducibility (if reported) | Summary of reference |
| --- | --- | --- | --- | --- | --- |
| Hietpas *et al.* (2013) | 6 or 7 | 568 | 30 | $R^2 = 0.95$ for full replicates | Beneficial single substitutions in Hsp90 of yeast under altered environments |
| Jiang *et al.* (2013) | 6 or 7 | 568 | 34 | $R^2 = 0.96$ for full replicates | Interaction of expression and single substitutions on DFE of yeast Hsp90 |
| Puchta *et al.* (2016) | 5-7 | $\approx 60\,000$ | 545.7 | $R^2 = 0.85$ | Network of epistatic interactions in yeast snoRNA |
| Roscoe *et al.* (2013) | 6 | 1 530 | 21 | $R^2 = 0.93$ for full replicates | Functional biophysics of single substitutions in ubiquitin of yeast |
| Hietpas *et al.* (2011) | 3 or 7 | 568 | 26 | $R^2 = 0.82$ for full replicates | DFE of single substitutions for a short region of Hsp90 in yeast |
| Bank *et al.* (2015) | 5 | 1 015 | 21.6 | credibility intervals (Fig. 6B) | DFE of epistatic substitutions in Hsp90 of yeast |
| Wu *et al.* (2013) | 3 | >400 | >0.002 | n/a | Compensatory single substitutions for neuraminidase mutant of influenza |
| Li *et al.* (2016) | 2 | 65 537 | 685.5 | $R^2 = 0.997$ mean across 15 pairs of biological replicates | Fitness landscape of a tRNA gene in yeast |
| Jiang *et al.* (2015) | 2 | 475 | 20.5 | $R^2 = 0.92$ for full replicates | Biophysics of single substitutions in influenza neuraminidase with antiviral |
| Roscoe and Bolon (2014) | 2 | 1 617 | 30 | $R^2 = 0.96$ for full replicates | Biophysics of single ubiquitin substitutions on E1 activity and yeast fitness |
| Melnikov *et al.* (2014) | 2 | 4 993 | 33.2 | $R^2 = 0.90$ for full replicates | Biophysics of single substitutions in APH(3')II in *E. coli* with antibiotics |
| Kim *et al.* (2013) | 2 | 29 708 | 90.2 | n/a | Biophysics of single substitutions in yeast Deg1 protein degradation signal |
| Melamed *et al.* (2013) | 2 | 110 745 | 186.5 | n/a | Biophysics of single and multiple substitutions in Pab1 RRM of yeast |
| Klesmith *et al.* (2015) | 2 | 9 219 | 5.8 | Reproducibility plot | Biophysics of levoglucosan consumption rate in *E. coli* |

## Model and Methods

### Experimental Setup

We consider an experiment assessing the fitness of *K* mutants that are labeled by *i* ∈ {1,2,…, *K*}. Each mutant is present in the initial library at population size *C_i_* and grows exponentially at constant rate *r_i_*. Consequently, the number of mutants of type *i* at time *t* is given by *N_i_*(*t*) = *C_i_* exp{*r_i_t*}. For convenience, we measure time in hours. Growth rates can easily be rescaled to (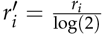), where *r_l_^′^* denotes the the growth rate per generation. At each sampling time point **t** = (*t*_1_ = 0,*t*_2_, …,*t_τ_*) from a multinomial distribution with parameters *D* (sequencing depth) and **p**(*t*) = (*p*_1_(*t*), p_2_(*t*),…, *P_K_*(*t*)), where (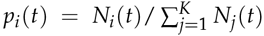) is the relative frequency of mutant *i* in the population at time *t*. Accordingly, *τ* and *t_τ_* denote the number of samples and the duration of the experiment, respectively. Note that for notational convenience, we will omit the subscript in *t* to denote any element in **t**. For illustrative purposes, we will present our results under the assumption that *T* equally spaced time points are sampled, such that **t** = (0,1,…, *T* − 1), and in particular *τ* = *T* and *t_τ_* = *T* − 1. Note that, with this definition,increasing the number of sampling time points (*T*) increases the actual numbers of samples taken (*τ*) *and* the duration of the experiment (*t_τ_*). The separate effects of *τ* and *t_τ_* will be discussed subsequently.

Furthermore, let **n**(*t*) = (*n*_1_(*t*),*n*_2_(*t*),…,*n_K_*(*t*)) denote the random vector of the number of sequencing reads sampled at time *t*. Without loss of generality, we denote the wild-type reference (or any chosen reference type) by *i* = 1 and set its growth rate to 1 (i.e., *r*_1_ = 1). Thus, mutant growth rates will be measured relative to that of the wild type. Accordingly, the selection coefficient of mutanti with respect to the wild type is given by, *s_i_* = *r_i_* −*r_1_*.

Estimators for the selection coefficients *s_i_* are then obtained from linear regression, based on log ratios of the number of sequencing reads *n_i_* (*t*) over the different sampling time points (but see Bank *et al.* 2014, for a Bayesian Markov chain Monte-Carlo approach). The corresponding linear model can then be written as

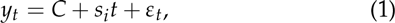

where *y_t_* is the (transformed) observation variable, *C* is a constant (i.e., the intercept) and *ϵ_t_* denotes the regression residual.

In the following, we derive an estimator that uses the log ratios of the number of reads of mutant i over the number of reads of the wild type as dependent variables in a linear regression. We call this method the *wild-type approach* (WT). In Supporting Information B we derive and analyze an alternative selection coefficient estimator that is based on log ratios of the number of mutant reads with respect to the total number of sequencing reads and which we call the *total approach* (TOT). This estimator has previously been used for detecting outliers within the experimental setup considered in Bank *et al.* (2014).

### Estimation of Selection Coefficients Ŝ_WT_

Ultimately, we want to calculate the mean of the log ratios of the number of sequencing reads for mutant *i* over the number of wild-type sequencing reads, (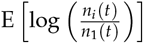). By noting that *n_i_*(*t*) is binomially distributed (for every mutant *i* ∈ {1,2,…, *K*}) and using the Delta method (for derivation see Supporting Information A; see also Hurt 1976; Casella and Berger 2002), we derive

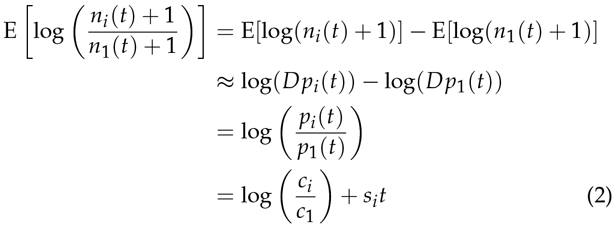

such that an estimator *Ŝ*_WT,*i*_ for *S_i_* can be obtained by applying the ordinary least squares (OLS) method on the linear regression model

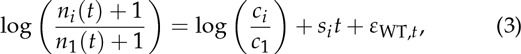

where *ɛ*_WT,*t*_ denotes the regression residual using the WT approach (as opposed to the TOT approach; see also Supporting Information B). Note that the additive term within the logarithm ensures that the logarithm is always well-defined and was added solely for mathematical convenience.

**Table 2.** Summary of notation and definitions.

|  |  |
| --- | --- |
| $K$ | number of mutants |
| $D$ | total number of reads per sampling time point (sequencing depth) |
| $c_i$ | initial population size of mutant $i$ |
| $r_i$ | (exponential) growth rate of mutant $i$ |
| $N_i(t)$ | number of mutants of type $i$ at time $t$ |
| $\mathbf{n}(t) = (n_1(t), n_2(t), \dots, n_K(t))$ | vector of the observed number of sequencing reads sampled at time $t$ |
| $p_i(t)$ | relative frequency of mutant $i$ at time $t$ |
| $\tau$ | number of samples taken |
| $t_\tau$ | duration of the experiment |
| $T$ | number of sampling time points ($\tau = T, t_\tau = T - 1$) |
| $s_i = r_i - r_1$ | selection coefficient of mutant $i$ |

### Simulation of Time-Sampled Deep Sequencing data

In order to validate analytical results, we simulated time-sampled deep sequencing data (implemented in C++; available upon request). We assumed that mutant libraries were created perfectly, such that the initial population size q was identical for all mutants and, accordingly, 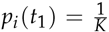 for all *i* = 1,2,…, *K*. Selection coefficients were independently drawn from a normal distribution with mean 0 and standard deviation 0.1. To test the robustness of these assumptions we performed additional simulations where initial population sizes were drawn from a log-normal distribution (i.e., (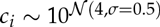) reflecting empirical distributions of inferred initial population sizes. Furthermore, selection coefficients were also drawn from a mixture distribution

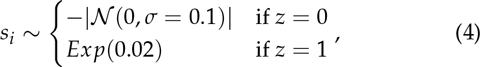

where *Z* ~ Bernoulli(0.7) (Fig. SI D_1). For a given number of sampling time points *T* and sequencing depth *D*, the number of mutant sequencing reads (*n*_1_(*t*), *n*_2_(*t*),…,*n*_*K*_(*t*)) was drawn from a multinomial distribution with parameters *D* and **p**(*t*) for each sampling point. Selection coefficient estimates (*Ŝ_i_*)*i* = 2,…, *K* were then obtained by fitting the linear model by means of OLS. Finally, the accuracy of the parameter estimates was assessed by computing the mean squared error (MSE),

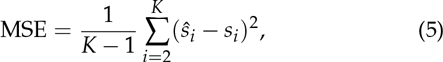

and the deviation (DEV)

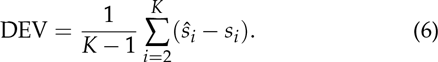

Note that we have omitted the hat over the MSE and DEV for notational convenience. If not stated otherwise, statistics were calculated over 1,000 simulated experiments for each set of parameters.

## Results and Discussion

The aim of this paper is to provide a statistical framework for *a priori* optimization of the experimental setup for future DMS studies. As such, our primary interest lies in the quantification of the MSE and its dependence on the experimental setup. We first deduce analytical approximations for the variance and the MSE of the estimators for the selection coefficient and compare these with simulated data. We then derive approximate formulas for the length of the confidence interval of the estimates and the mean absolute error (MAE), which can be used to assess the expected precision of the estimates. For each of these steps, we discuss the consequences of relaxing some of the above assumptions along with potential extensions of the model. Finally, we apply our statistical framework to experimental evolution data of 568 engineered mutations from Hsp90 in *Saccharomyces cerevisiae*, and show that our model indeed captures the most prevalent source of error (i.e., error from sampling).

### Approximation of the Mean Squared Error

Generally, the MSE of an estimator 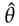 (for parameter *θ* is given by

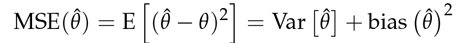

(see section 7.3.1 of Casella and Berger 2002). Since E [*ɛ*_WT_ = 0 (i.e., the mean of the regression residual is zero, implying that *Ŝ*_WT,*i*_ is an unbiased estimator; Fig. SI D_2), it is sufficient to analyze Var [*Ŝ*_WT,*i*_] to asses MSE (*Ŝ*_WT,*i*_). For ease of notation, and since all results in the main text are derived using the wild-type approach, we will omit the wT index from here on. Taking the variance of equation (3) implies

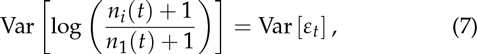

which, by applying the Delta method (see Supporting Information A) and using equation (S5) together with equations (S4) and (S6) can be approximated by

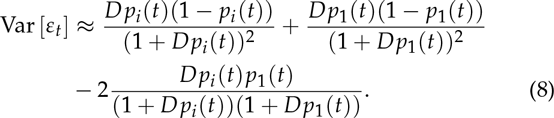

Note that the residuals are heteroscedastic (i.e., their variance is time-dependent; the relative mutant frequencies *p_i_*(*t*) change during the course of the experiment). Hence, there is no general closed form expression of the variance of *Ŝ_i_*. However, by making the simplifying assumption of homoscedasticity (i.e., *p_i_*(*t*) ≈ *p_i_*(*t*_1_) and *p*_1_(*t*) ≈ *p*_1_(*t*_1_) for all *t*), we obtain

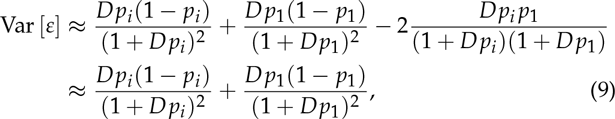

where the dependence on time has been dropped for ease of notation. Note that omitting the covariance term implicitly assumes that the number of mutants *K* is sufficiently large (i.e., *p*_i_ and *p*_1_ are small). we discuss the effect of assuming homoscedastic error terms below. Equation (9) shows that Vars [*ɛ*] decreases monotonically with increasing sequencing depth and increasing relative proportions of the wild-type and focal mutants.

Using existing theory on variances of slope coefficients in a linear regression framework with homoscedastic error terms (e.g., see section 11.3.2 Casella and Berger 2002), the variance of the selection coefficient estimate is given by

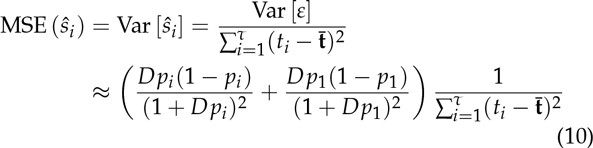

which is our first main result.

Using that sampling times are assumed to be equally spaced, equation (10) can further be rewritten as

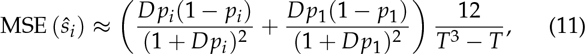

which shows that the MSE decreases cubically with the number of time points *T* (Fig. 1). Thus, sampling additional time points (i.e., taking more samples *and* extending the duration of the experiment) drastically increases the precision of the measurement.

**Figure 1.**
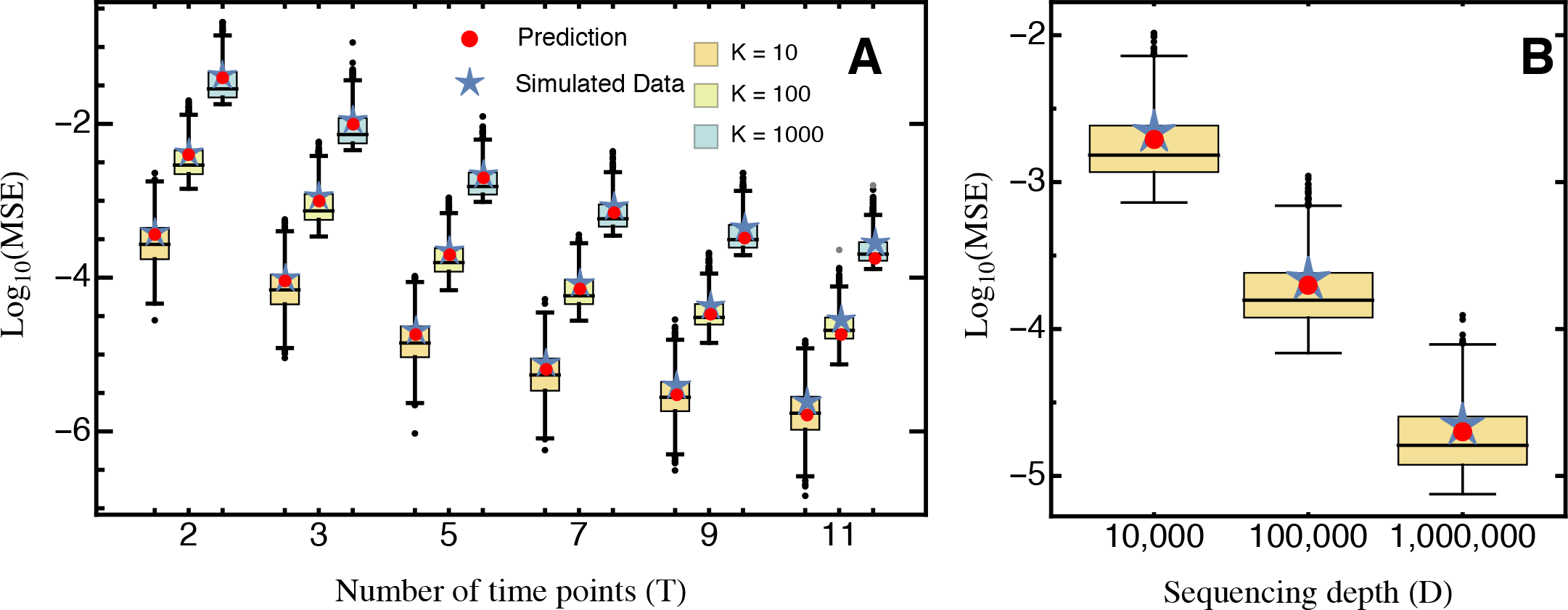
Comparison of the predicted mean squared error (eq. 10; red) and the average mean squared error (blue star), obtained from 1,000 simulated data sets for (A) different numbers of sampling time points *T* and mutants *K*, for sequencing depth *D* = 100,000, and for (B) different sequencing depth *D*, with fixed *T* = 5, *K* = 100. Boxes represent the interquartile range (i.e., the 50% C.I.), whiskers extend to the highest/lowest data point within the box ± 1.5 times the interquartile range, and black and gray circles represent close and far outliers, respectively. Results are presented on log-scale.

Our approximation generally performs very well across the entire parameter space. Although we assumed that the relative abundance of all mutants remains roughly constant with time (i.e., neglecting that error terms are heteroscedastic), the (small) absolute error of our approximation remains constant across time points (Fig. SI D_3A). Deviations from homoscedasticity increase as more and later time points are sampled, as shown by the relative error (Fig. SI D_3B). This is also reflected by the deviation between the predicted MSE and the true average MSE obtained from the simulated data (Fig. 1). For realistic experimental durations however, compared to the experimental error (due to sampling) this approximation error is negligible.

***Uneven sampling schemes***. To obtain a closed formula for the decay in the measurement error with the number of time samples *T* (eq. 11), we assumed equally spaced sampling times. The observed decay remains cubic relative to the number of time points also when samples are not taken at equally spaced time points. Furthermore, equation (10) informs about the optimal sampling scheme to use to minimize measurement error: for fixed sequencing depth and number of mutants, the MSE is minimized when the sum of squared deviations of the sampling times from their mean is maximized. In other words, to minimize the measurement error one should sample in two sampling blocks one at the beginning and another at the end of the experiment instead of sampling throughout the experiment, or, if time and resources allow, create full two-time-point replicates (e.g., *t* = (0,1,5,6) is better than *t* = (0,2,4,6); see also the interactive demonstration tool provided online).

***Duration and sampling density of the experiment***. Equation (11) implies that the MSE decreases cubically when both more samples are taken and the duration of the experiment is extended. However, extending the experiment indefinitely is impossible, both because of experimental constraints and because secondary mutations will begin to affect the measurement. Hence, the possible duration of an experiment under a given condition may be a (fixed) short time *t_τ_* (e.g., less than 20 yeast generations for *EMPIRIC).* To separate the effects of taking more samples *τ* from those of extending the duration of the experiment *t_τ_*-which are combined in T in the normal model setup (see Model and Methods)-equation (11) can be re-written as

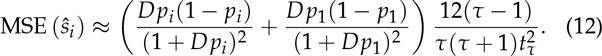

Thus, when the duration of the experiment *t_τ_* is held constant, measurement error decays linearly as *τ* (i.e., the number of sampling points) increases. Conversely, when extending the duration of experiment, the MSE decreases quadratically. This result suggests that the experimental duration should always be maximized under the constraints that mutants grow exponentially and population size is much lower than the carrying capacity. How long both of these assumptions are met depends on each individual mutant’s selection coefficient (or growth rate) and its initial frequency. Accordingly, there is no universal ‘optimal’ duration of the experiment. For example, the frequency of strongly deleterious mutations in the population generally decreases quickly, such that the phase where they show strict exponential growth is short and does not span the entire duration of the experiment. Furthermore, mutations might be lost from the population before the experiment is completed. Thus, when sampling two time points that extend over a long experimental time, growth rate estimates for strongly deleterious mutations can be substantially overestimated (see also Contribution of additional error: Data application).

Conversely, for mutations with small (i.e., wild-type-like) selection coefficients, increasing the duration of the experiment considerably improves the precision of the estimates. Specifically, to infer deviations from the wild-type’s growth rate the (expected) log ratio of the number of mutant sequencing reads over the number of wild-type sequencing reads (i.e., the ratio of relative frequencies between mutant and wild-type abundance) need to change consistently with time (i.e., either increase or decrease; eq. 2). However, changes in the log ratios will be small if the duration of the experiment is short, and even if there are slight shifts, sequencing depth *D* needs to be large enough such that they are not washed-out by sampling.

Thus, beyond the linear improvement on the MSE that comes with increasing *τ*, sampling more time points can be an efficient strategy to capture the entire range of selection coefficients (i.e., strongly deleterious and wild-type like mutants). Specifically, sampling in two blocks (one at the beginning and another at the end of the experiment as suggested above) would allow using different *t_τ_* depending on the underlying selection coefficient, which could be determined by a bootstrap leave-p-out cross validation approach (for details see Contribution of additional error: Data application). For example, the first sampling block could be used for strongly deleterious mutations, whereas all sampled time points could be used for the remaining mutations, reducing error due to overestimation of strongly-deleterious selection coefficients and increasing statistical power to detect differences to wild-type like growth rates.

***Library design and the number of mutants***.Increasing the number of mutants *K* reduces the number of sequencing reads per mutant and hence *p_i_*;, which explains the approximately linear increase of the MSE with *K* (Fig. 1). Crucially, we assumed that the initial mutant library was balanced, such that all mutants were initially present at equal frequencies. In practice this is hardly ever the case and previous analyses have shown that initial mutant abundances instead follow a log-normal distribution (Bank *et al.* 2014). Taking this into account, we find that unbalanced mutant libraries, as expected from equation (9), introduce an error due to the higher variance terms resulting from the generally lower *p_i_*; (Fig. SI D_5). This error can be avoided by using the estimated relative mutant abundance, 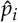, in equation (9) (Fig. SI D_5).

The additional-though practically inevitable-error introduced by variance in mutant abundance indicates that library preparation is an important first step for obtaining precise estimates. In fact, equation (9) suggests that the measurement precision increases with the relative abundance of the wild type (such that the second term in eq. 9 decreases). However, this results in a trade-off because increasing wild-type abundance results in a decrease of the abundance of all other mutants, which leads to an increase of the first term in equation (9). Assuming that increasing the relative abundance of the wild type reduces the relative abundance of all mutants equally (i.e., *p_i_* =*p_j_* for all *i, j* ɛ {2,3,…, *K*}), we find that precision is maximized by increasing the wild-type abundance by a factor proportional to 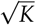o (analytical result not shown; Fig. 2). This way, the MSE can be reduced by 50% as compared to the MSE with equal proportions of all mutants. Most importantly however, if wild-type abundance is low (i.e., 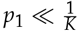), the error increases substantially (i.e., more than 10-fold; see inset in Fig. 2 A).

***Sequencing depth and its fluctuations***. The MSE decreases approximately linearly with the sequencing depth *D* (Fig. 1), because the number of reads per mutant increases. As long as *D* is independent of the number of mutants *K* in the actual experiment, it can simply be treated as a rescaling parameter; hence, qualitative results are independent of the actual choice of *D*. Similarly, the variance of the estimated MSE decreases approximately quadratically with sequencing depth and increases quadratically as the number of mutants increases (Fig. 1).

Although we here treat the sequencing depth D as a constant parameter, it will in practice vary between sampling time points. Thus, D should rather be interpreted as the (expected) average sequencing depth taken over all time points. In particular, compared to a fixed sequencing depth, variance in *D* introduces an additional source of error (due to increased heteroscedasticity), although deviations from the predicted to the observed mean MSE remain roughly identical (Fig SI D_6). Our model can also account for other forms of sampling. For example, if the sample taken from the bulk competition is known to be smaller than the sequencing depth, its size should be used as *D* in the precision estimates.

**Figure 2.**
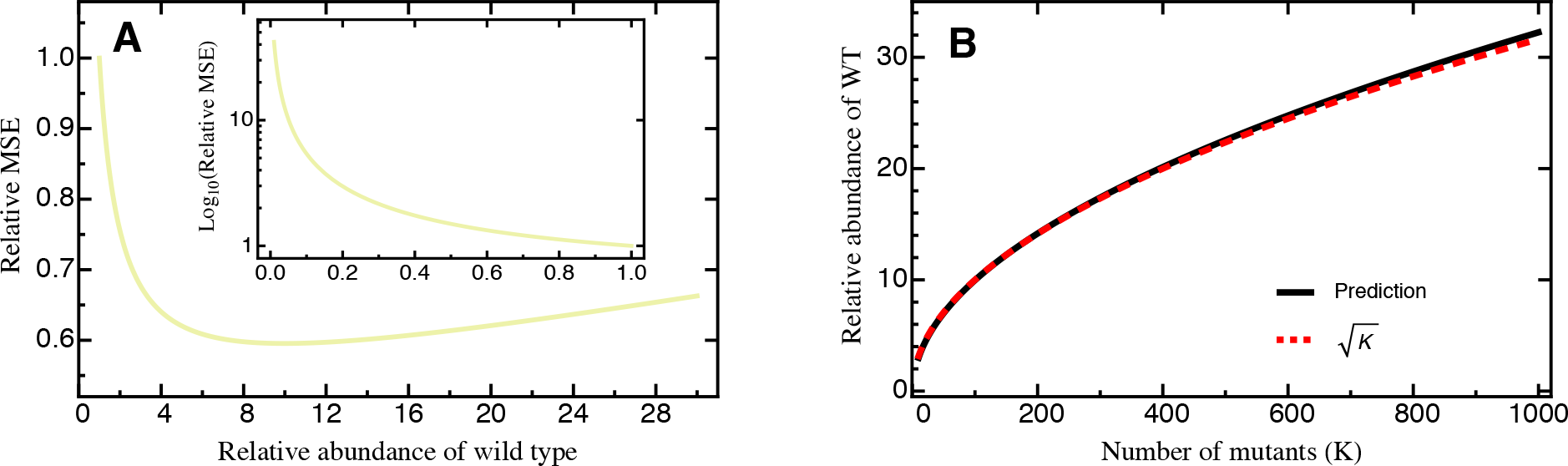
**A** The relative MSE as a function of the relative abundance of the wild type, i.e., 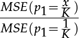, for *K* = 100. The inset shows results for 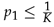,where the y-axis has been put on log-scale. The abundance of all other (except the wild type) is assumed to scale proportionally. **B** The relative wild-type abundance which minimizes the MSE as a function of the number of mutants *K*. Explicit formula (as given by the black line) is not shown due to complexity, but can well be approximated by 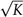. Either prediction is based on equation (10). Other parameters: *D* = 100,000.

***Shape of the underlying DFE***. Our results remain qualitatively unchanged when selection coefficients are drawn from differently shaped DFEs. The assumed normally distributed DFE corresponds to theoretical expectations derived from Fisher’s geometric model (assuming that the number of traits under selection is large; Martin and Lenormand 2006a; Tenaillon 2014). DFEs inferred from experimental evolution studies, however, are typically characterized by an approximately exponential tail of beneficial mutations and a heavier tail of deleterious mutations (Eyre-Walker and Keightley 2007; Bank *et al.* 2014) that roughly follows a (displaced) gamma distribution (Martin and Lenormand 2006a; Keightley and Eyre-Walker 2010). To account for this expected excess of deleterious mutations in the DFE (reviewed by Bataillon and Bailey 2014), we used a mixture distribution that resulted in a highly skewed DFE. For this, beneficial mutations (s > 0) were drawn from an exponential distribution and deleterious mutations were given by the absolute value drawn from a Gaussian distribution (Fig. SI D_1; see Methods for details). Even with this highly skewed DFE, we did not find changes to either the MSE (Fig. SI D_4) or the deviation (Fig. 3), indicating that our results are robust across a range of realistic DFEs.

***An alternative normalization***. In Supporting Information B, we analyze and discuss an alternative estimation approach based on the log ratios of the number of mutant reads over the sequencing depth *D* (as opposed to a single reference/wild type) that was proposed in Bank *et al.* (2014) and called the “total” (TOT) approach. Although the TOT approach can improve results for very noisy data (i.e., if *T* or *D* are small; Figs. SI B_1, SI B_2, SI B_3, SI B_4), its estimates are generally biased. The bias increases with the number of time points and overrides the smaller variance in residuals (see eqs. 9 and S8). Thus, application of the TOT approach is only recommended under special circumstances, e.g. under the suspicion of outlier measurements in the wild type (as in the case of Bank *et al.* 2014).

### Confidence Intervals, Precision and Hypothesis Testing

One way of quantifying the precision of the estimated selection coefficient is obtained using Jensen’s inequality (see section 6.6 of Williams 1991), which yields an upper bound for the mean absolute error (MAE)

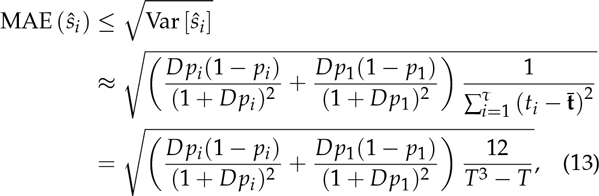

where, in the last line, we have again assumed that sampling times are equally spaced. Thus, the MAE is simply the square root of the MSE.

Alternatively, using central limit theorem arguments (Rice 1995), it can be shown that for a fixed mutant *i* the estimated selection coefficient **Ŝ_i_** asymptotically follows a normal distribution (Figs. 3, SI D_2). The upper and lower bound of the (1 – *α*)-confidence interval with significance level *α* for *s_i_* are then given by

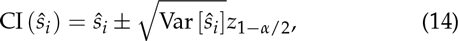

where *z*_1 − *α*/2_ denotes the (1 − *α*/2)-quantile of the standard normal distribution. The length of the (1 − *α*)-confidence interval, *L*_(1 − *α*)_, can be used as an intuitive *a priori* measure for the precision of the estimated selection coefficient. Formally, let 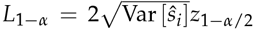 denote the length of the (1 – α)-confidence interval. Setting *α* = 0.05 and using equation (10), we obtain the approximation

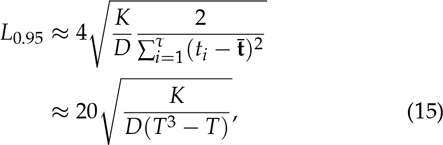

where we assumed Z_0.975_ ≈ 2. Equation (15) shows that the sequencing depth *D* and the number of mutants *K* are inversely proportional. Similarly to equation (10), the number of time points *T* enters cubically.

**Figure 3.**
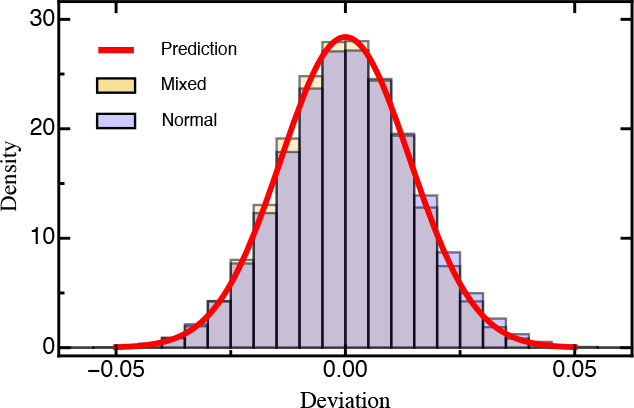
Histogram of the deviation (eq. 6) between the estimated and true selection coefficients either drawn from a normal distribution or a mixture distribution (for details see Model and Methods) based on 1,000 simulated data sets each. The red line is the prediction based on equation 10. Other parameters: *T* = 5, *D* = 100,000, *K* = 100.

Furthermore, equation (14) can be used to define the upper and lower bounds of the region of rejection of a two-sided Z-test with, for instance, null hypothesis *s_i_* = 0 (or more generally any other null hypothesis *s_i_* = *θ*). The Z-statistic is then given by

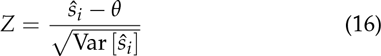

(see chapter 8 in Sprinthall 2014). This statistic can be applied to existing data to test whether a mutant has an effect different from the wild type. In addition, we can use this statistic to determine the maximum achievable statistical resolution of a planned experiment.

### Optimization of Experimental Design

Equation (10) suggests that the measurement error modelled here could in theory be eliminated entirely by sampling (infinitely) many time points. In practice, the attainable resolution of the experiment is also limited by technical constraints imposed by the experimental details and by sequencing error, and by the available manpower and budget. To further improve the experimental design taking the latter two factors into account, we can integrate our approach into an optimization problem using a cost function *C_α, β, C_tτ_, K_*(*D, t_τ_, τ*). As an example, we define

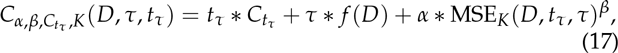

where *C_t_τ__* denote the personnel costs over the duration of the experiment, *f*(*D*) denotes the sequencing costs per sampled time point, and *α* and *β* scale the associated error costs given by equation (12) (Boyd and Vandenberghe 2004). The optimization problem is solved by minimizing

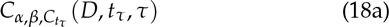

under constraints

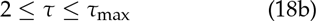

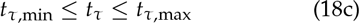

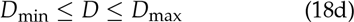

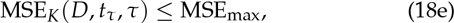

which yields the maximum tolerable error MSE_max_ while minimizing the total experimental costs. An illustrative example is given in Supporting Information C.

### Contribution of Additional Error: Data Application

An important limitation of our model is that it does not consider additional sources of experimental error. Therefore, any results presented here should be interpreted as upper limits of the attainable precision. In particular, sequencing error (dependent on the sequencing platform and protocol used) is expected to affect the precision of measurements. However, if the additional error is non-systematic (i.e., random), it will not change the results qualitatively, but solely add an additional variance to the measurement.

To assess the influence of additional error sources to the validity of our statistical framework, we re-analyzed a data set of 568 engineered mutations from Hsp90 in Saccharomyces cerevisiae grown in standard laboratory conditions (i.e., 30°C; for details see Bank et al. 2014). We estimated the initial population size and the selection coefficient for each mutant using the linear-regression framework discussed here. With respect to the experimental parameters (i.e., number and location of sampling points, sequencing depth) and our proposed model, we simulated 1,000 bootstrap data sets. We assessed the accuracy of our selection coefficient estimates by calculating the MSE between the selection coefficient estimates obtained from the bootstrap data sets and those obtained from the experimental data, which serve as a reference for the “true” (but unknown) selection coefficient. To quantify the effect of the number of sampling time points, we used a leave-p-out cross-validation approach, successively dropping sampling time points (Geisser 1993).

**Figure 4.**
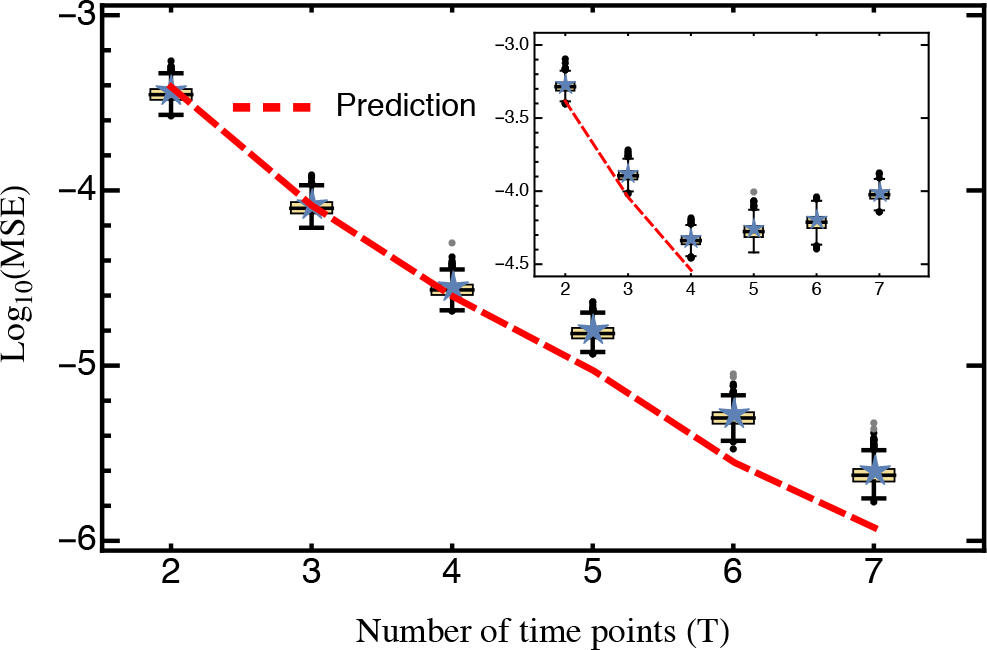
Comparison of the predicted mean squared error (eq. 10; red) against the average mean squared error (blue star) obtained from 1,000 cross-validation data sets. Only mutants with an estimated selection coefficient larger than the intermediate between the estimated mean synonym and the estimated mean stop codon selection coefficient were considered. The inset shows the MSE calculated for all mutants. The MSE is presented on log-scale. Other parameters: **t** = (4.8,7.2,9.6,12,16.8,26.4,36), *D*_Figure_ = (474,931,636,257,873,827,1,513,392,424,182,443,739,452,326), *D*_Inset_ = (654,311,820,301,1,046,169,1,726,516,469,855,464,070,463,363), *K*_Figure_ = 400, *K*_Inset_ = 568.

For the complete data set, our prediction holds only when the number of time points considered is small. Conversely, with more than four time samples, the MSE even slightly increases with the number of sampling points (inset in Fig. 4). However, when strongly deleterious mutations (i.e., those with a selection coefficient closer to that of the average stop codon than to the wild type, see also Bank *et al.* 2014) are excluded from the analysis, the MSE is very well predicted by equation (10) for any number of time points (Fig. 4). Two model violations may well explain the observed pattern when deleterious mutations are included. Firstly, the frequency of strongly deleterious mutations in the population decreases quickly and do not show strictly exponential growth (Fig. S2 in Bank *et al.* 2014), especially for later time points. Secondly, these mutations might not be present in the population over the entire course of the experiment. Sequencing error will then create a spurious signal, feigning and extending their “presence”, thus biasing the results. The bootstrap approach utilized here could in principle be used to determine the time points that should be considered for the estimation of strongly deleterious mutations, and to generally test for model violations. Indeed, Figure 4 demonstrates that our model captures the most prevalent source of error (i.e., error from sampling) when strongly deleterious mutations are excluded.

## Conclusion

The advent of sophisticated biotechnological approaches on a single-mutation level, combined with the continual improvement and reduction in costs of sequencing, present us with an unprecedented opportunity to address long-standing questions about mutational effects and the shape of the distribution of fitness effects. An additional step towards optimizing results receives little attention: by systematically invoking statistical considerations ahead of empirical work, it is possible to quantify and maximize the attainable experimental power while avoiding unnecessary expenses, both regarding financial and human resources. Here, we present a thorough statistical analysis that results in several straightforward, general predictions and rules of thumb for the design of DMS studies, which can be applied directly to future experiments using a free interactive web tool provided online (https://evoldynamics.org/tools). We emphasize here three important and general rules that emerged from the analysis:

1. Increasing sequencing depth and the number of replicate experiments is good, but adding sampling points together with increasing the duration of the experiment is much better for accurate estimation of small-effect selection coefficients.
2. Preparation of a balanced library is the key to good results. The quality of selection coefficient estimates strongly depends on the abundance of the reference genotype: Always ensure that the frequency of the reference genotype is larger than 1/*K*-“less is a mess”.
3. Clustering sampling points at the beginning and the end of the experiment increases experimental power, and allows the efficient and precise assessment of the entire range of the distribution of fitness effects.
Although the statistical advice presented here is limited to experimental approaches that fulfill the requirements listed in the introduction and focuses on the error introduced through sampling, our work highlights the promises that lie in long-term collaborations between theoreticians and experimentalists as compared to the common practice of post-hoc statistical consultation.

## Acknowledgements

We thank Ivo Chelo, Ines Fragata, Isabel Gordo, Kristen Irwin, and Adamandia Kapopoulou for helpful discussion and comments on earlier versions of this manuscript. This project was funded by grants from the Swiss National Science Foundation (FNS) and a European Research Council (ERC) Starting Grant to
JDJ.

## Supplemental Information

### A. Derivation of the Delta method

In this Supporting Information we will briefly motivate and introduce the delta method and derive the equations in the main text. Consider a generic random variable *X* with finite second moment and smooth function *f*: ℝ → ℝ. If *f* is non-linear (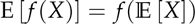) does in general no longer hold such that *f* and (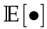) can no longer be interchanged. However, an approximate result can be obtained by Taylor-expanding *f* around (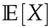) up to the second order such that

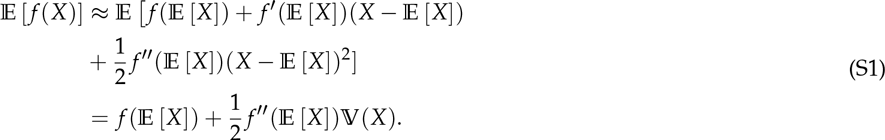

This approach is called the Delta-method or method of error propagation (Hurt 1976; Oehlert 1992; Casella and Berger 2002). Thus, for *X* ~ Bin(*D, p*) and *f*(*x*) = log(1 + *x*), the expectation of *f*(*X*) can be approximated as

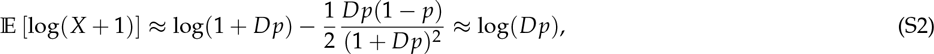

which is used in equation (2) in the main text. Note that the approximation induces a small error. However, for fixed *p* this distortion is of order *O*(*D*^−1^), which is generally negligible if *D* is large (i.e., when the sequencing depth is large).

Analogously, we can calculate the variance of *f*(*X*) as

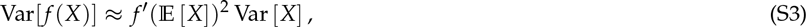

which again for *X* ~ Bin(*D, p*) and *f*(*x*) = log(1 + *x*) becomes

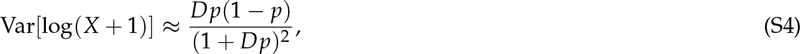

which is used in equation (9) and (10) in the main text.

Similarly, let *g*: ℝ → ℝ be a smooth function and *X*_1_, *X*_2_ denote two square integrable random variables. Taylor-expanding up to the first order and taking variances yields

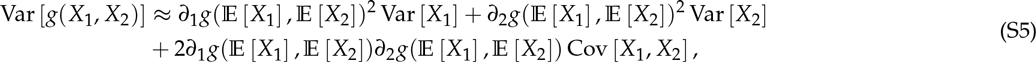

which is used for the derivation of equation (9) and (10) in the main text. Furthermore, if *x*_1_ and *x*_2_ denote two realizations of the same multinomial, the covariance between *x*_1_ and *x*_2_ are given by

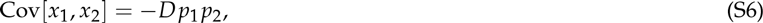

which is again used in equation (9) and (10) in the main text.

### B. The Total Approach

In this Supporting Information B we derive and analyze an alternative estimator of selection coefficient to the one proposed in the main text. Unlike the WT approach this estimator is based on the log-ratios of the number of mutant reads with respect to the *total* number of sequencing reads (i.e., sequencing depth) and which is thus called the *total approach* (TOT). This estimator has previously been used for detecting outliers in time-sampled deep-sequencing bulk completion data (Bank *et al.* 2014) and proved to be more robust than the WT approach. Analogous to the main text we will first analyse and discuss the statistical properties of selection coefficient estimator based on the TOT approach and then compare its performance to the WT approach. Finally, we will end by giving rough guidelines when to prefer one over the other approach.

***Statistical analysis of the TOT approach*** In contrast to the WT approach the TOT approach is based on the log-ratios of the number of mutant reads with respect to the sequencing depth *D*, such that a linear model can be written as

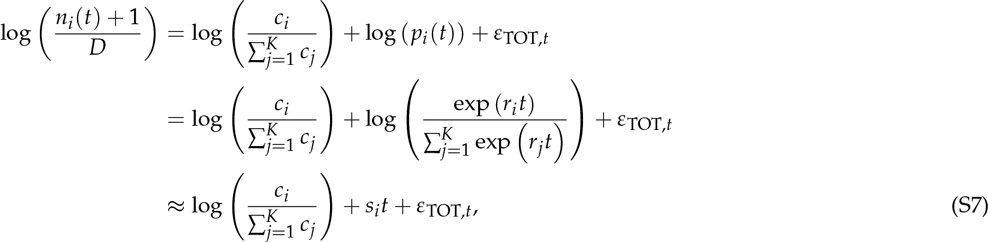

where the approximation assumes that *r*_j_ ≈ 1 for all mutants *K*.

Then, with a calculation analogous to the one in the main text, we obtain

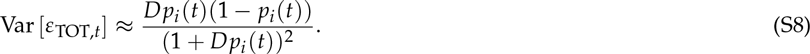

Again assuming that residuals are homoscedastic, i.e., that *p_i_*(*t*) ≈ *p_i_*(*t*_1_) for all *t*, we have

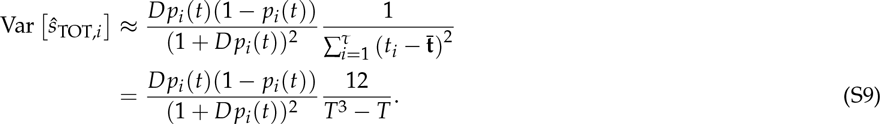

However, the approximation used in equation (S7) induces a systematic error, such that E[*ɛ*_TOT,*t*_] ≠ 0 meaning that ŝ_TOT,*i*_ is generally biased as can be seen from Figure SI B_1 and Figure SI B_2.

**Figure SI B_1.**
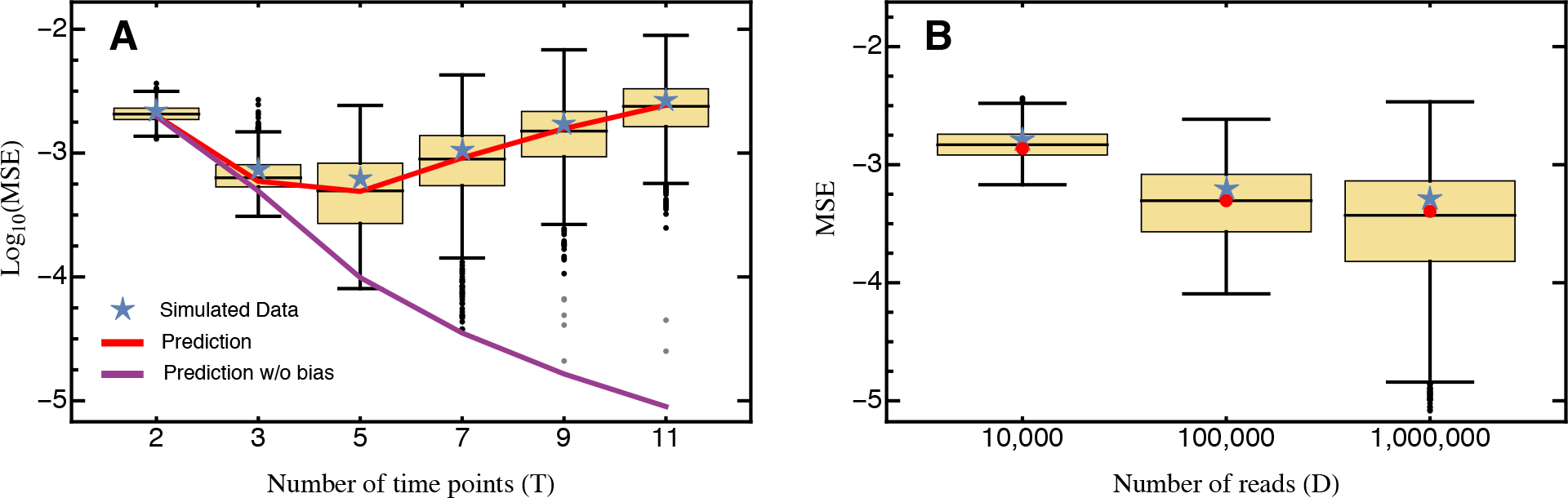
The average mean squared error (blue star) obtained from 1,000 simulated data sets for **A** different numbers of sampling time points *T* compared against the analytical prediction with (red) and without (purple) accounting for the estimation bias as given by equation S15 and S9, respectively. **B** for different numbers of sequencing reads *D* with *T* = 5. Boxes represent the interquartile range (i.e., the 50% C.I.), whiskers extend to the highest/lowest data point within the box ± 1.5 times the interquartile range, and black and gray circles represent close and far outliers, respectively. Results are presented on log-scale. Other parameters: *D* = 100,000, *K* = 100.

**Figure SI B_2.**
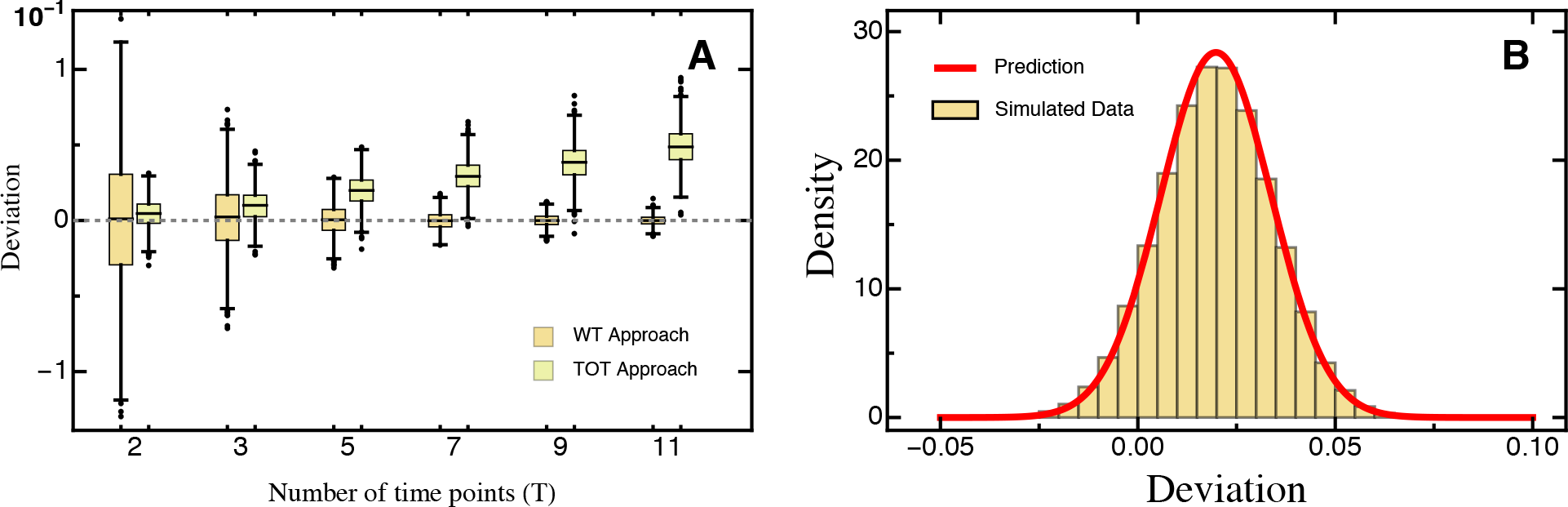
**A** Comparison of the deviation (eq. 6) of the true and estimated selection coefficient as obtained by the WT and TOT approach, calculated from 1,000 simulated data sets for different numbers of sampling time points *T*. **B** Histogram of the deviation (eq. 6) between the estimated and true selection coefficients based on 1,000 simulated data sets using the TOT approach with *T* = 5. The red line is a normal distribution centered around the empirical mean and variance given by the square root of equation (S9). Other parameters: *D* = 100,000, *K* = 100.

To quantify this bias, we will now derive an approximation based on the Delta method (see Supporting Information A). Rewriting equation (S7) we obtain

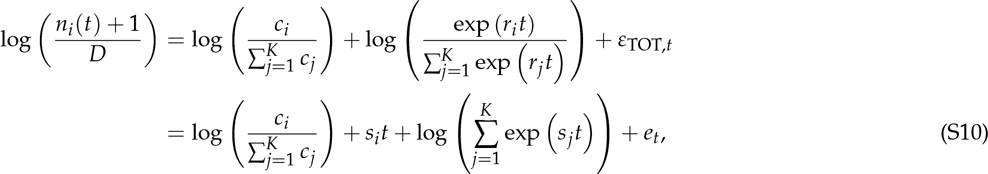

where (*e*_*t*)*t*=,…,*T*_ are independent with mean zero, and 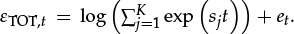. For ease of notation we will define (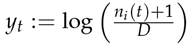) and 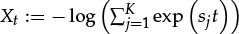, and use the fact that the slope coefficient of a simple linear regression can be expressed as

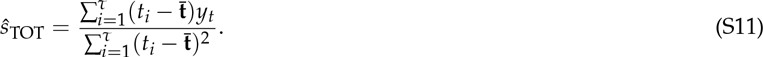

Defining (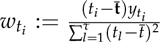), and using that (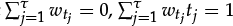), and E[*e_t_*] = 0, we obtain

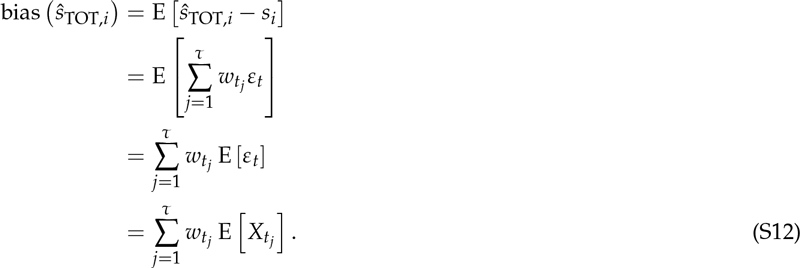

Note that in order to derive an explicit formula for the bias, the distribution of the (*s*_*i*)*i*=2,… *K*_ needs to be specified. In accordance with our simulation assumptions, we consider the case where the (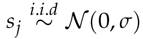). Hence, the random variables (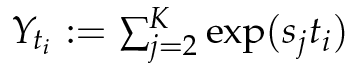)are

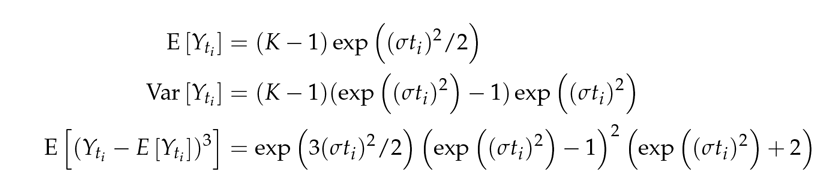

Since 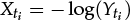), we have that Taylor-expanding up to the third order and taking expectations yields

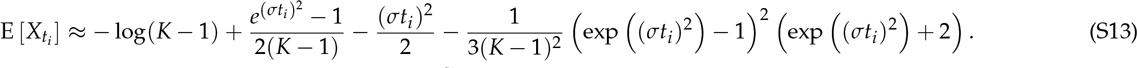

Finally, combining equations (S12) and (S13) and using that (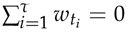) yields

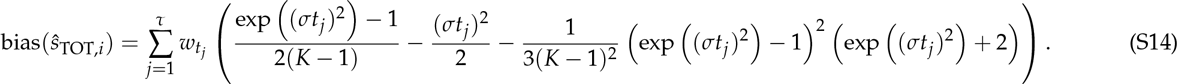

Note that the accuracy of the approximation strongly depends on *σ, τ* and *t_τ_* potentially because of the local validity implied by the Taylor approximation breaking down. In particular, for small *σ* and *T* (i.e., *σ* ≤ 0.1, *τ* ≤ 7 and *t_τ_* ≤ 7) yields a reasonable prediction of the bias (Figs. SI B_1, SI B_3).

Thus, by combining equations (S9) and (S14) we obtain

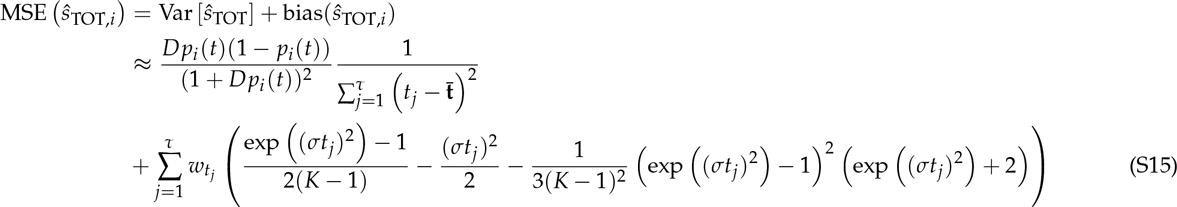

while comparing equations (9) and (S9) shows that Var [*ŝ*_TOT,*i*_] < Var [*ŝ*wT,*i*] (with identical *T, D* and *p_i_*), the bias clearly limits the use of the TOT approach. In particular, the bias is strongest when *T* (and in particular *t_τ_*) is large and/or only a few mutants were considered (i.e., *K* is low relative to *D*). Similarly, increasing *D* does not improve the mean MSE by much (Fig. SI B_1).

**Figure SI B_3.**
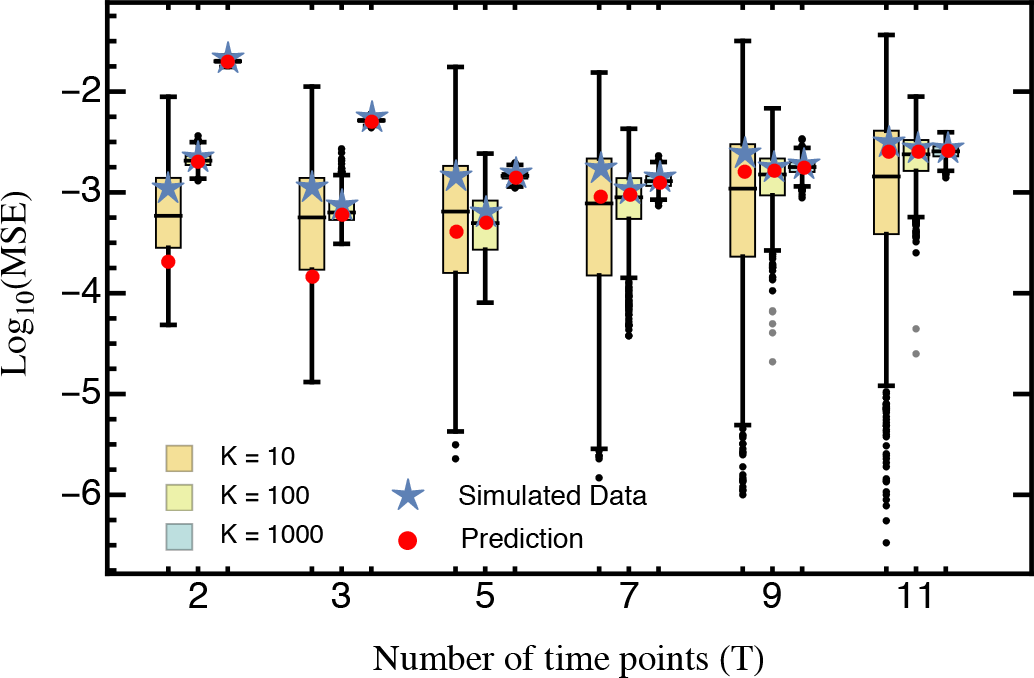
Comparison of the predicted mean squared error (eq. S15; red) against the average mean squared error (blue star) obtained from 1,000 simulated data sets for different numbers of sampling time points *T* and mutants *K*. Other parameters: *D* = 100,000.

***Comparison of the WT and TOT approach*** Comparing the TOT approach to the wT approach (Fig. SI B_4) shows that the former only outperforms the latter when stochastic forces (induced by the multinomial sampling and only a few sampling time points) are large (i.e., when *D* and *T* are small, and *K* is big)-in other words whenever there were problems with obtaining the data. This also shows up by the generally reduced variance of the MSE when increasing *K* or decreasing *D* (Figs. SI B_1, SI B_3, SI B_4). This is due to the fact that calculating the log-ratios with respect to the sequencing depth *D* introduces a “saturation effect” such that the log-ratios are non-linear in *t* as if the mutants no longer grow exponentially (comparable to deceleration and saturation phase described in Hall *et al.* 2014). In line with our observations (Figs. SI B_1, SI B_3, SI B_4), this effect is strongest if *K* is small and/or *T* is large (and the duration of the experiment is long), i.e., if *p_i_* becomes large such that the log-ratios saturate. Accordingly, the bias also strongly depends on the variance of the DFE and thus on the environment. If mutants can generally grow faster, i.e., if the variance in the DFE is high, the log-ratios will start to become saturated even earlier (i.e., with smaller *T*). The overall effect is that selection coefficient estimates will be less extreme, i.e., large positive selection coefficients (strongly beneficial mutants) will be underestimated whereas large negative selection coefficients (strongly deleterious mutants) will be overestimated. Still, when the wild type is systematically misestimated (due to some non-random error in the experiment) or if the the wild type is rare (see also Fig 2) the TOT approach might outperform the WT approach. Furthermore, when applied for detecting outliers-where this approach has been proposed initially and proved to be more robust (Bank et al. 2014)-only the TOT approach is able to detect potential outliers in the wild type. In particular, when outliers in the wild type remain undetected they will introduce a bias in the estimated selection coefficients for all other mutants. Accordingly, calculating the log-ratios with respect to the number of sequencing reads of wild-type like mutants could make use of the advantages of the WT and TOT approach, i.e., reducing the variance of the estimator without introducing a bias, which is however beyond the scope of this manuscript.

**Figure SI B_4.**
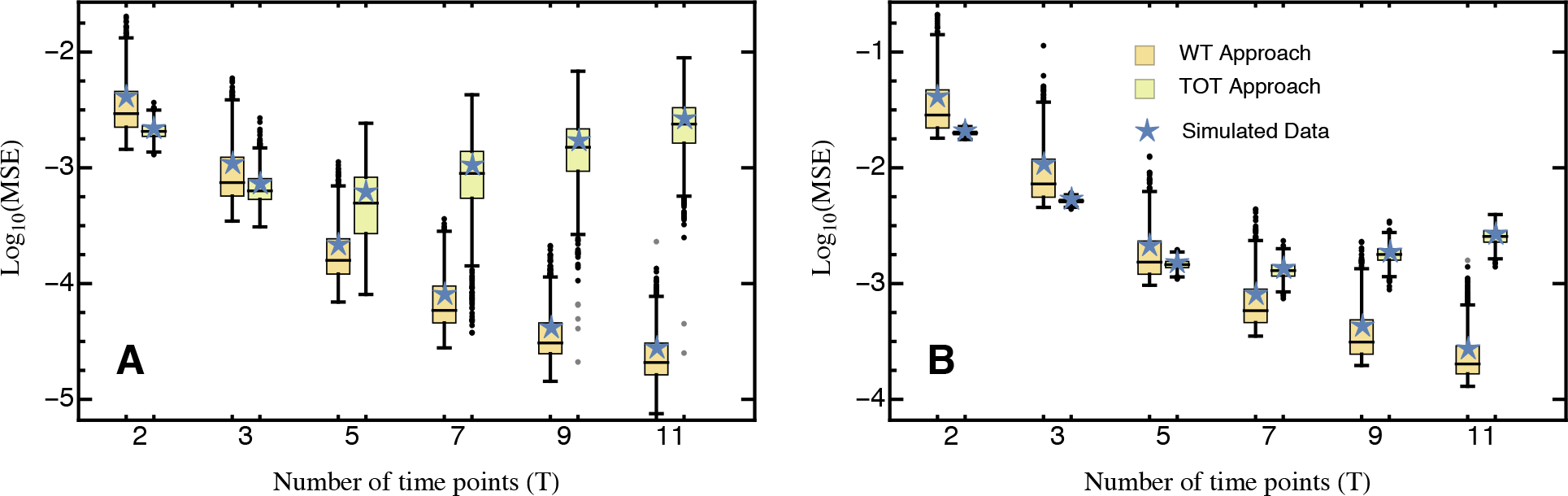
Comparison of the mean squared error obtained under the WT and TOT approach calculated from 1,000 simulated data sets for different numbers of sampling time points *T*. **A** *K* = 100 **B** *K* = 1000. Note the differences in scale. Results are presented on log-scale. Other parameters: *D* = 100,000.

### C. Optimization Example

To illustrate the potential use of the optimization approach, we parameterized equation (17) as

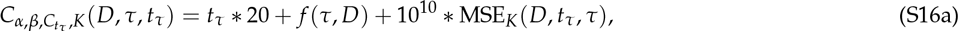

where

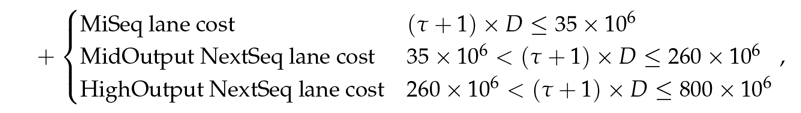

which is inspired by the experimental protocol described in Hietpas *et al.* (2012). The first term in equation (S16a) denotes personnel costs as given by product of the duration of the experiment and the median per hour salary of a post-doc researcher in the US (according to payscale.com).

Estimated order-of-magnitude costs for the procedures given in *f* (*τ*, *D*) were either estimated from personal lab experience or taken from the Georgia Genomics Facility as

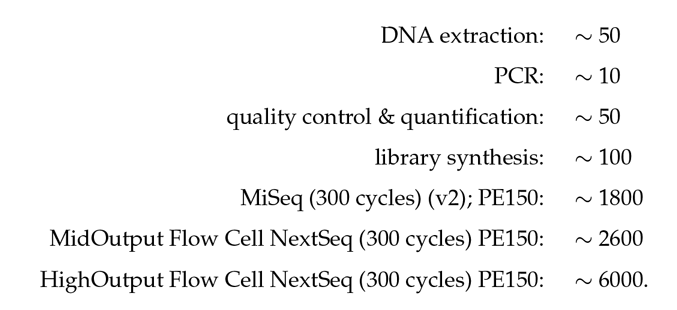

Note that following Hietpas *et al.* (2012), an additional “processing analysis” sequencing run is done per experiment on the wild-type sequence in order to determine the misread analysis, which is why the total number of reads per experiment is (*τ* + 1) × *D*. Furthermore we have chosen *α* = 10^10^ and *β* = 1 such that a higher MSE induces additional costs reflecting that stochasticity in the experiment can lead to larger than expected errors (that are highly penalized with this parametrization, putting an emphasise on the quality of the estimate rather than on experimental costs). Thus, the lower the MSE the more likely it is that the desired minimal precision MSE_max_ is reached.

Constraints are chosen arbitrarily since they depend on the experimental setup and are given as

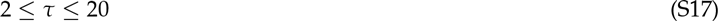

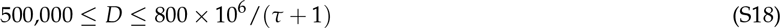

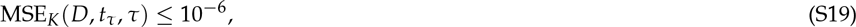

reflecting that over the course of the experiment at most 20 samples can be taken, and that at least 500,000 but up to 800 × 10^6^/ (*τ* + 1) sequencing reads can be obtained. Furthermore, we assume that the experiment is designed to infer the growth rates of *K* = 10,000 mutants that evolve for a fixed experimental duration *t_τ_* = 20, and the desired minimal precision (given by MSE_max_) is 10^−6^ (corresponding to MAE = 10^−3^; eq. 13). In addition, we for simplicity assume that sampling time points are equally spaced such that the sampling interval is given by *t_τ_* / *τ*.

Figure SI C_1 shows that under the given constraints costs (Min *C*_*α, β*, *C_t τ_*, *K*_(*D, τ, t_τ_*)) sharply decrease and are minimized for *τ*(*D, τ, t_τ_*)) sharply decrease and are minimized for *τ* = 7 with a sequencing depth *D* = 1 × 10^8^, but start to increase again when more time points are sequenced. Pure monetary costs, however, monotonically increase in t. Thus, if budgetary constraints were emphasized (e.g., with *α* = 0), it would be optimal to only sample twice at the beginning and the end of the experiment. As argued above, however, sampling additional time points (in particular more than two) can improve capturing the entire range of selection coefficients.

Note that in principle any cost function could be used (also incorporating additional cost terms). Furthermore, sequencing costs and progress in sequencing technology are inherently dynamic factors, such that any parametrization of *C*_*α, β*, *C_t τ_*, *K*_(*D, τ, t_τ_*)) sharply decrease and are minimized for *τ*(*D, τ*) becomes outdated and unrealistic immediately. However, as demonstrated by the above example: The experimental guidelines derived from this study can be used as an auxiliary tool for assessing the a priori measurement error, and, when specifying experimental costs, to design cost-and time-efficient experiments. Specifically, by providing a free, web-based interface, cost functions can be tailored to the specific experimental setup and thus our results can readily be used to design efficient and statistically robust experiments.

**Figure SI C_1.**
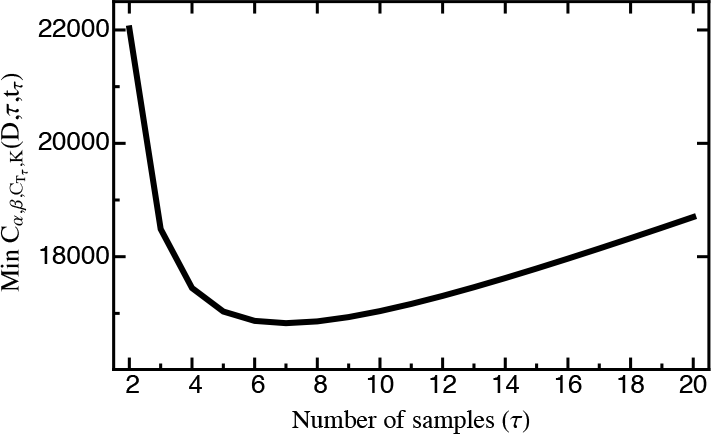
Min*C*_*α, β*, *C_t τ_*, *K*_(*D, τ, t_τ_*)as parameterized by eq. S16a for different τ. Costs are minimized for *τ* = 7. Note however, that pure monetary costs monotonically increase in *τ*.

### D. Supporting Figures

**Figure SI D_1.**
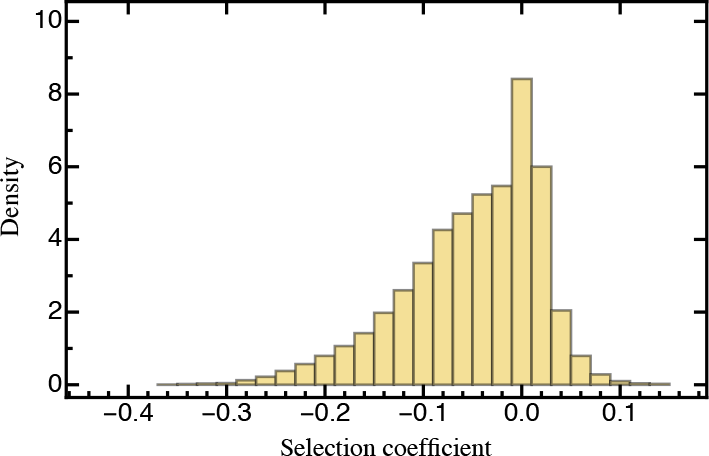
Histogram showing the distribution of fitness effects obtained from 10,000 draws from a mixture distribution that serves as an alternative to the normal distribution that is otherwise assumed throughout the manuscript. For details see Model and Methods.

**Figure SI D_2.**
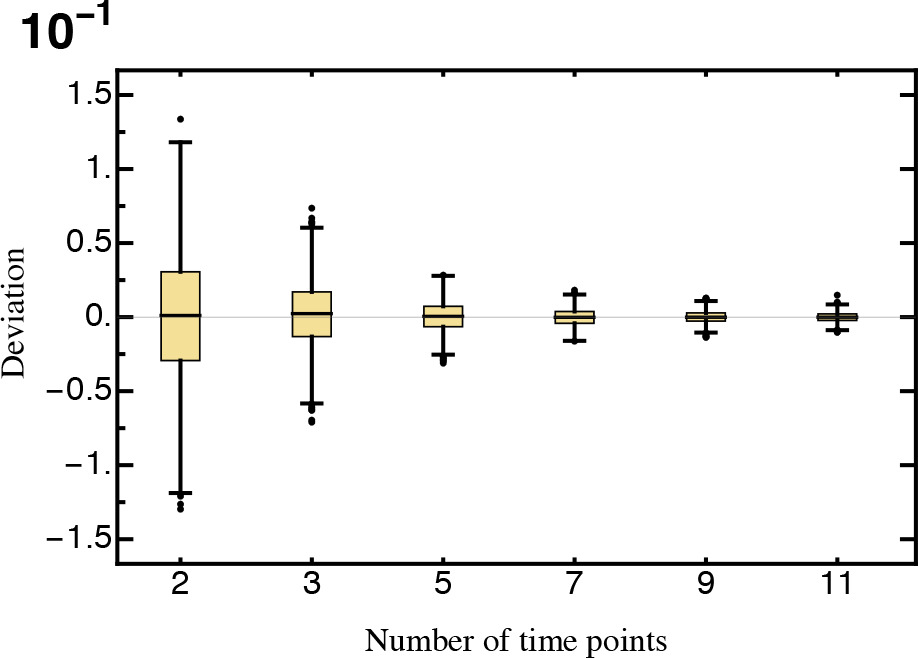
The deviation (eq. 6) of the estimated selection coefficient from the true selection coefficient obtained from 1,000 simulated data sets for different numbers of sampling time points. Other parameters: *D* = 100,000, *K* = 100.

**Figure SI D_3.**
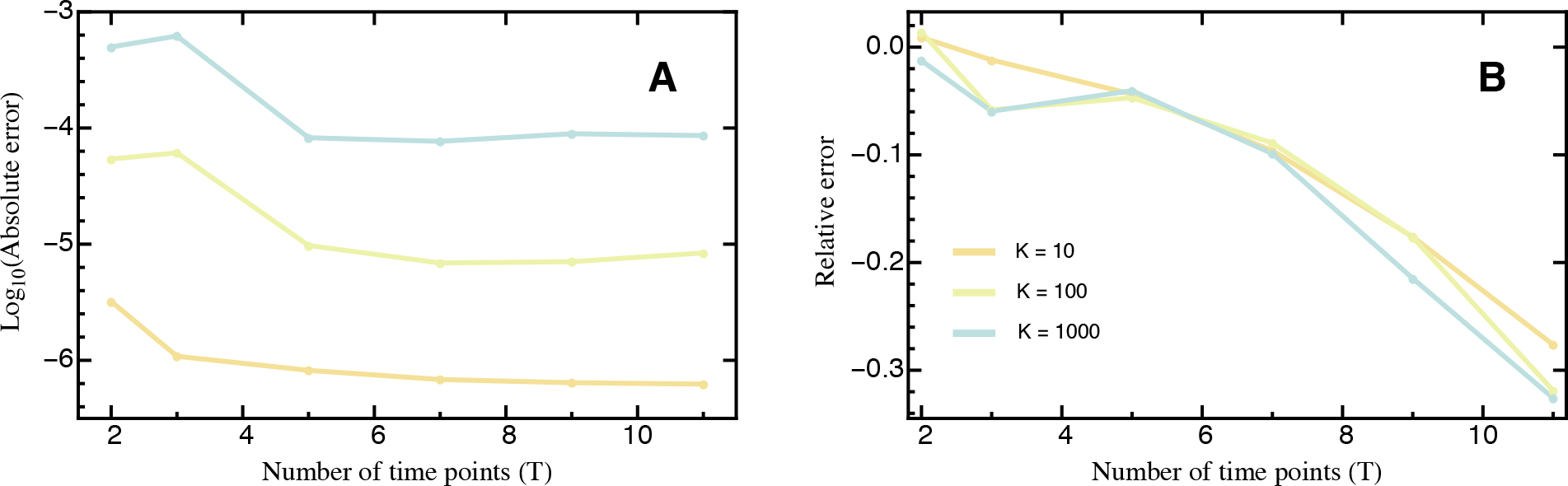
The absolute (**A**) and the relative (**B**) error calculated from 1,000 simulated data sets for different numbers of sampling time points *T* and mutants *K*. Results in **A** are presented on log-scale. Other parameters: *D* = 100,000.

**Figure SI D_4.**
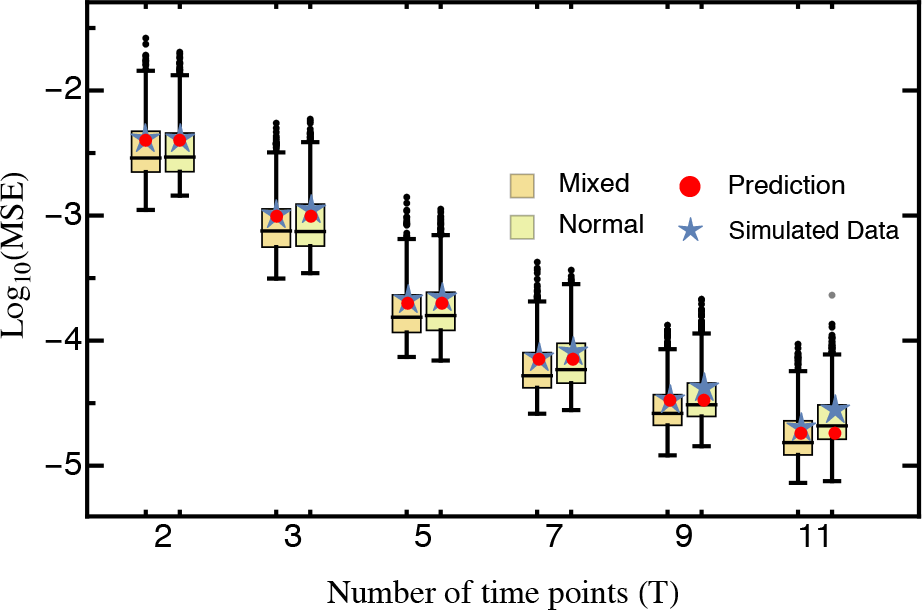
Comparison of the predicted mean squared error (eq. 10; red) against the average mean squared error (blue star) obtained from 1,000 simulated data sets for different numbers of sampling time points. Results are presented on log-scale. Otherparameters: *D* = 100,000, *K* = 100.

**Figure SI D_5.**
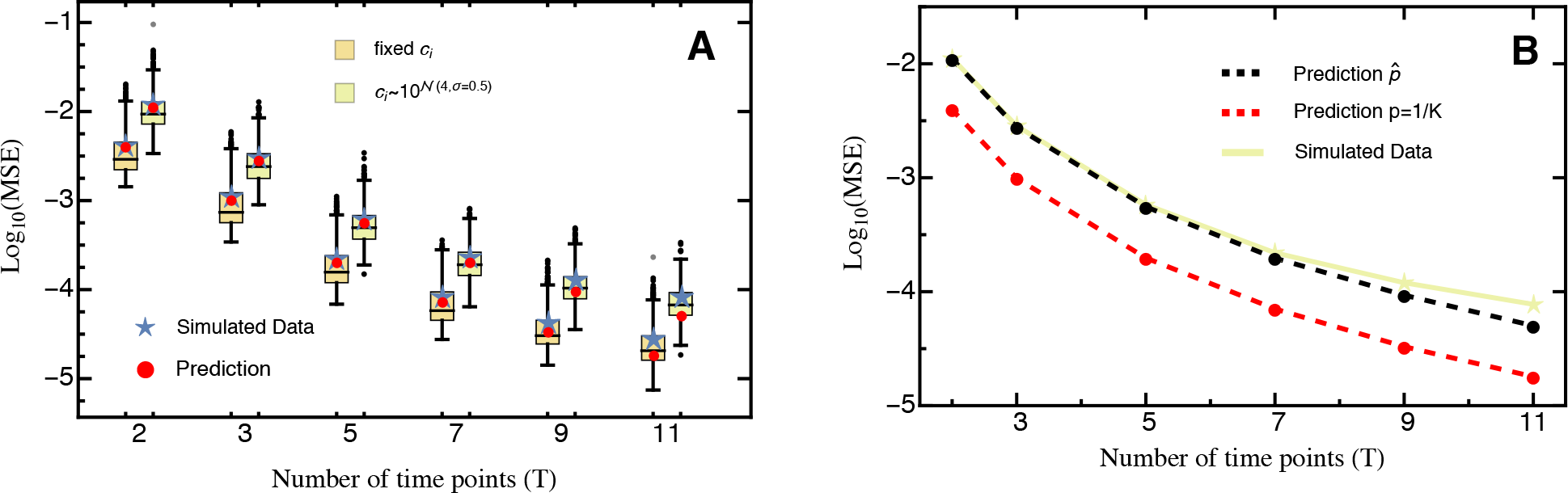
The mean squared error obtained from 1,000 simulated data sets with a fixed initial population size *c_i_* compared to that where *c_i_* has been drawn from a log-normal distribution. In **A** the prediction (eq. 10; red) is based on (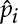), i.e., the estimated relative abundance of mutant *i* such that it can be referenced against the empirical average MSE (blue stars). **B** compares the empirical average MSE (solid line) against the predicted MSE with a balanced mutant library (i.e., *p* = 1/*K*; black) and an uneven mutant library (i.e., (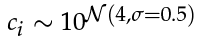); red). Either prediction is based on equation (10). Note the differences in scale. Results are presented on log-scale. Other parameters: *D* = 100,000.

**Figure SI D_6.**
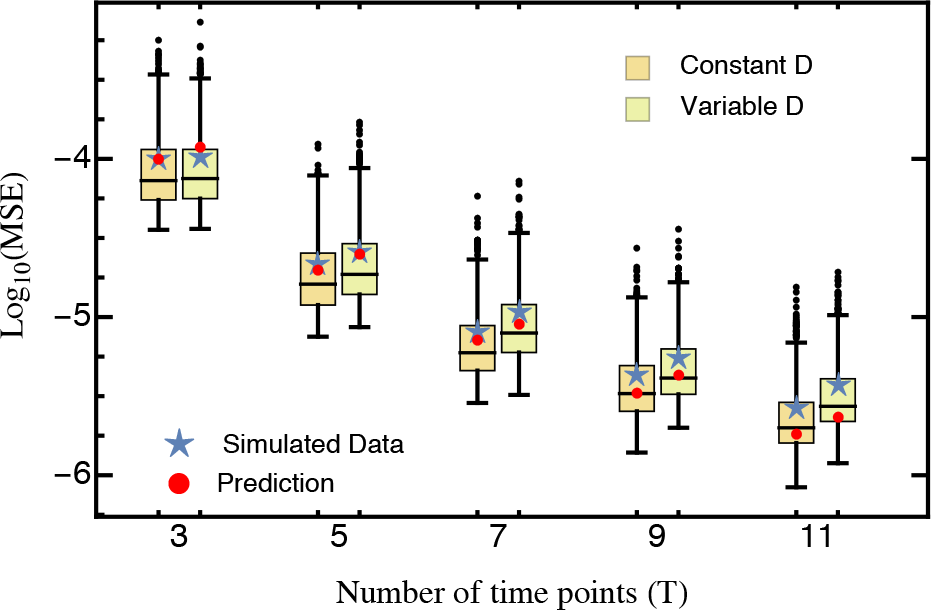
The mean squared error obtained from 1,000 simulated data sets with a fixed sequencing depth *D* compared to that where *D* alternates between 1,000,000 and 500,000 for each time point. The predicted mean squared error (eq. 10; red) for the varying sequencing depth is calculated based on average sequencing depth across time points. The empirical average MSE is given by the blue stars. Results are presented on log-scale. Other parameters: *K* = 100.

**Figure SI D_7.**
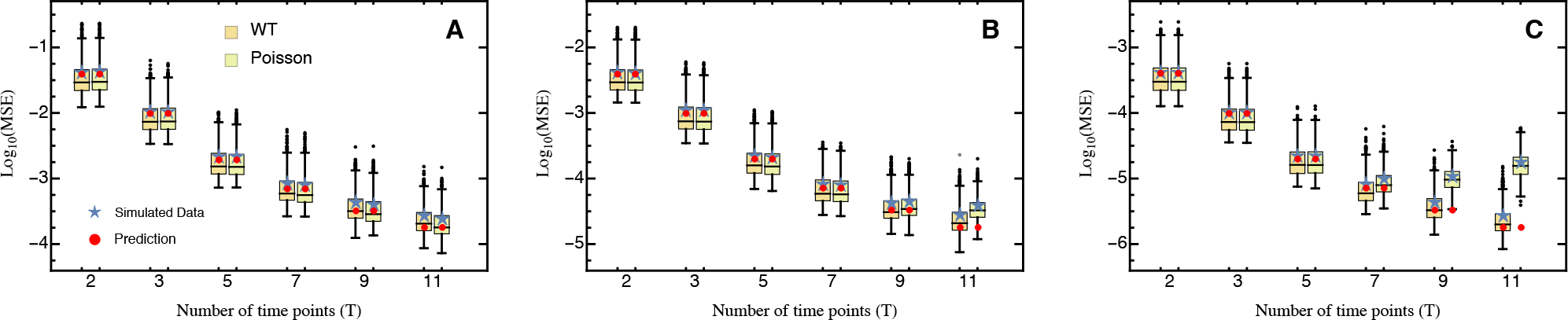
Comparison of the predicted mean squared error (eq. 10; red) against the average mean squared error (blue star) obtained from 1,000 simulated data sets using the WT-approach and a Poisson regression approach of the sequencing counts against time with sequencing depth **A** *D* = 10,000 **B** *D* = 100,000 **C** *D* = 1,000,000. Results are shown on log-scale. Note the differences in y-axis range. Other parameters: *K* = 100.

## Literature Cited

Bank, C., R. T. Hietpas, J. D. Jensen, and D. N. Bolon, 2015 A systematic survey of an intragenic epistatic landscape. Mol. Biol. Evol. 32: 229–238.

Bank, C., R. T. Hietpas, A. Wong, D. N. Bolon, and J. D. Jensen, 2014 A bayesian mcmc approach to assess the complete distribution of fitness effects of new mutations: Uncovering the potential for adaptive walks in challenging environments. Genetics 196: 841–852.

Bataillon, T. and S. F. Bailey, 2014 Effects of new mutations on fitness: insights from models and data. Ann. N. Y. Acad. Sci. 1320: 76–92.

Bernet, G. P. and S. F. Elena, 2015 Distribution of mutational fitness effects and of epistasis in the 5′ untranslated region of a plant rna virus. BMC Evol. Biol. 15: 1–13.

Boucher, J. I., D. N. A. Bolon, and D. S. Tawfik, 2016 Quantifying and understanding the fitness effects of protein mutations: Laboratory versus nature. Protein Science pp. n/a–n/a.

Boyd, S. and L. Vandenberghe, 2004 Convex optimization. Cambridge University Press.

Casella, G. and R. L. Berger, 2002 Statistical inference, volume 2. Duxbury Pacific Grove, CA.

Charlesworth, D., B. Charlesworth, and M. T. Morgan, 1995 The pattern of neutral molecular variation under the background selection model. Genetics 141: 1619–1632.

Chen, F., S. M. Pruett-Miller, Y. Huang, M. Gjoka, K. Duda, J. Taunton, T. N. Collingwood, M. Frodin, and G. D. Davis, 2011 High-frequency genome editing using ssdna oligonucleotides with zinc-finger nucleases. Nat. Meth. 8: 753–755.

Connallon, T. and A. G. Clark, 2015 The distribution of fitness effects in an uncertain world. Evolution 69: 1610–1618.

Eyre-Walker, A., 2010 Genetic architecture of a complex trait and its implications for fitness and genome-wide association studies. Proc. Natl. Acad. Sci USA 107: 1752–1756.

Eyre-Walker, A. and P. D. Keightley, 2007 The distribution of fitness effects of new mutations. Nat. Rev. Genet. 8: 610–618.

Firnberg, E., J. W. Labonte, J. J. Gray, and M. Ostermeier, 2014 A comprehensive, high-resolution map of a gene’s fitness landscape. Molecular Biology and Evolution 31: 1581–1592.

Fowler,D. M., C. L. Araya, S. J. Fleishman, E. H. Kellogg, J. J. Stephany, D. Baker, and S. Fields, 2010 High-resolution mapping of protein sequence-function relationships. Nat. Meth. 7: 741–746.

Fowler, D. M. and S. Fields, 2014 Deep mutational scanning: a new style of protein science. Nature Methods 11: 801–807.

Frenkel, E. M., B. H. Good, and M. M. Desai, 2014 The fates of mutant lineages and the distribution of fitness effects of beneficial mutations in laboratory budding yeast populations. Genetics 196: 1217–1226.

Geisser, S., 1993 Predictive Inference. Chapman & Hall/CRC Monographs on Statistics & Applied Probability, Taylor & Francis.

Gerrish, P. J. and R. E. Lenski, 1998 The fate of competing beneficial mutations in an asexual population. Genetica 102/103: 127–144.

Gillespie, J. H., 1983 A simple stochastic gene substitution model. Theor. Pop. Biol. 23: 202–215.

Gordo, I. and P. R. A. Campos, 2013 Evolution of clonal populations approaching a fitness peak. Biol. Lett. 9: rsbl20120239.

Hall, B. G., H. Acar, A. Nandipati, and M. Barlow, 2014 Growth rates made easy. Mol. Biol. Evol. 31: 232–238.

Halligan, D. L. and P. D. Keightley, 2010 Spontaneous mutation accumulation studies in evolutionary genetics. Annu. Rev. Ecol. Evol. Syst. 40: 151–172.

Hietpas, R., B. Roscoe, L. Jiang, and D. N. A. Bolon, 2012 Fitness analyses of all possible point mutations for regions of genes in yeast. Nat. Protoc. 7: 1382–1396.

Hietpas, R. T., C. Bank, J. D. Jensen, and D. N. A. Bolon, 2013 Shifting fitness landscapes in response to altered environments. Evolution 67: 3512–3522.

Hietpas, R. T., J. D. Jensen, and D. N. A. Bolon, 2011 Experimental illumination of a fitness landscape. Proc. Natl. Acad. Sci. USA 108: 7896–7901.

Hurt, J., 1976 Asymptotic expansions of functions of statistics. Aplikace matematiky 21: 444–456.

Imhof, M. and C. Schlotterer, 2001 Fitness effects of advantageous mutations in evolving Escherichia coli populations. Proc. Natl. Acad. Sci. USA 98: 1113–1117.

Jacquier, H., A. Birgy, H. Le Nagard, Y. Mechulam, E. Schmitt, J. Glodt, B. Bercot, E. Petit, J. Poulain, G. Barnaud, P.-A. Gros, and O. Tenaillon, 2013 Capturing the mutational landscape of the beta-lactamase tem-1. Proceedings of the National Academy of Sciences 110: 13067–13072.

Jensen, J. D., K. R. Thornton, and P. Andolfatto, 2008 An approximate bayesian estimator suggests strong, recurrent selective sweeps in drosophila. PLoS Genet. 4: e1000198.

Jiang, L., P. Liu, C. Bank, N. Renzette, K. Prachanronarong, L. S. Yilmaz, D. R. Caffrey, K. B. Zeldovich, C. A. Schiffer, T. F. Kowalik, J. D. Jensen, R. W. Finberg, J. P. Wang,, and D. N. Bolon, 2015 A balance between inhibitor binding and substrate processing confers influenza drug resistance. J. Mol. Biol. 428: 538–553.

Jiang, L., P. Mishra, R. T. Hietpas, K. B. Zeldovich, and D. N. A. Bolon, 2013 Latent effects of hsp90 mutants revealed at reduced expression levels. PLoS Genet. 9: e1003600.

Jinek, M., K. Chylinski, I. Fonfara, M. Hauer, J. A. Doudna, and E. Charpentier, 2012 A programmable dual-rna-guided dna endonuclease in adaptive bacterial immunity. Science 337: 816–821.

Joung, J. K. and J. D. Sander, 2013 Talens: a widely applicable technology for targeted genome editing. Nat. Rev. Mol. Cell. Biol. 14: 49–55.

Keightley, P. D. and A. Eyre-Walker, 2010 What can we learn about the distribution of fitness effects of new mutations from dna sequence data? Phil. Trans. R. Soc. B 365: 1187–1193.

Kim, I., C. R. Miller, D. L. Young, and S. Fields, 2013 High-throughput analysis of in vivo protein stability. Mol. Cell. Pro-teomics 12: 3370–3378.

Kimura, M., 1979 Model of effectively neutral mutations in which selective constraint is incorporated. Proc. Natl. Acad. Sci. USA 76: 3440–3444.

Klesmith, J. R., J.-P. Bacik, R. Michalczyk, and T. A. Whitehead, 2015 Comprehensive sequence-flux mapping of a levoglu-cosan utilization pathway in e. coli. ACS Synthetic Biology 4: 1235–1243, PMID: 26369947.

Kowalsky, C. A., J. R. Klesmith, J. A. Stapleton, V. Kelly, N. Re-ichkitzer, and T. A. Whitehead, 2015 High-resolution sequence-function mapping of full-length proteins. PLoS ONE 10: 1–23.

Li, C., W. Qian, C. J. Maclean, and J. Zhang, 2016 The fitness landscape of a trna gene. Science 352: 837–840.

Martin, G. and T. Lenormand, 2006a A general multivariate extension of Fisher’s geometrical model and the distribution of mutation fitness effects across species. Evolution 60: 893907.

Martin, G. and T. Lenormand, 2006b The fitness effect of mutations in stressful environments: a survey in the light of fitness landscape models. Evolution 60: 2413–2427.

Melamed, D., D. L. Young, C. E. Gamble, C. R. Miller, and S. Fields, 2013 Deep mutational scanning of an rrm domain of the saccharomyces cerevisiae poly(a)-binding protein. RNA 19: 1537–1551.

Melnikov, A., P. Rogov, L. Wang, A. Gnirke, and T. S. Mikkelsen, 2014 Comprehensive mutational scanning of a kinase in vivo reveals substrate-dependent fitness landscapes. Nucleic Acids Res. 42: e112.

Oehlert, G. W., 1992 A note on the delta method. Am. Stat. 46: 27–29.

Ohta, T., 1977 Molecular evolution and polymorphism. National Institute of Genetics, Mishima, Japan.

Ohta, T., 1992 The nearly neutral theory of molecular evolution. Annu. Rev. Ecol. Syst. 23: 263–286.

Olson, C. A., N. C. Wu, and R. Sun, 2014 A comprehensive biophysical description of pairwise epistasis throughout an entire protein domain. Current Biology 24: 2643–2651.

Orr, H. A., 1998 The population genetics of adaptation: the distribution of factors fixed during adaptive evolution. Evolution 52: 935–949.

Orr, H. A., 2005a The genetic theory of adaptation: a brief history. Nat. Rev. Genet. 6: 119–127.

Orr, H. A., 2005b Theories of adaptation: what they do and don’t say. Genetica 123: 3–13.

Orr, H. A., 2009 Fitness and its role in evolutionary genetics. Nat. Rev. Genet. 10: 531–539.

Puchta, O., B. Cseke, H. Czaja, D. Tollervey, G. Sanguinetti, and G. Kudla, 2016 Network of epistatic interactions within a yeast snorna. Science 352: 840–844.

Rice, D. P., B. H. Good, and M. M. Desai, 2015 The evolutionarily stable distribution of fitness effects. Genetics 200: 321–329.

Rice, J., 1995 Mathematical Statistics and Data Analysis. Number Bd. 1 in Duxbury advanced series, Duxbury Press.

Rokyta, D. R., P. Joyce, S. B. Caudle, and H. A. Caudle, 2005 An empirical test of the mutational landscape model of adaptation using a single-stranded dna virus. Nat. Genet. 37: 441–444.

Roscoe, B. P. and D. N. A. Bolon, 2014 Systematic exploration of ubiquitin sequence, e1 activation efficiency, and experimental fitness in yeast. J. Mol. Biol. 426: 2854–2870.

Roscoe, B. P., K. M. Thayer, K. B. Zeldovich, D. Fushman, and D. N. A. Bolon, 2013 Analyses of the effects of all ubiquitin point mutants on yeast growth rate. J. Mol. Biol. 425: 1363–1377.

Rozen, D. E., J. A. G. M. de Visser, and P. J. Gerrish, 2002 Fitnesseffects of fixed beneficial mutations in microbial populations. Curr. Biol. 12: 1040–1045.

Sarkisyan, K. S., D. A. Bolotin, M. V. Meer, D. R. Usmanova, A. S. Mishin, G. V. Sharonov, D. N. Ivankov, N. G. Bozhanova, M. S. Baranov, O. Soylemez, N. S. Bogatyreva, P. K. Vlasov, E. S. Egorov, M. D. Logacheva, A. S. Kondrashov, D. M. Chudakov, E. V. Putintseva, I. Z. Mamedov, D. S. Tawfik, K. A. Lukyanov, and F. A. Kondrashov, 2016 Local fitness landscape of the green fluorescent protein. Nature 533: 397–401.

Sawyer, S. A., R. J. Kulathinal, C. D. Bustamante, and D. L. Hartl, 2003 Bayesian analysis suggests that most amino acid replacements in drosophila are driven by positive selection. Journal of Molecular Evolution 57: S154–S164.

Schneider, A., B. Charlesworth, A. Eyre-Walker, and P. D. Keight-ley, 2011 A method for inferring the rate of occurrence and fitness effects of advantageous mutations. Genetics 189: 1427–1437.

Sousa, A., S. Magalhaes, and I. Gordo, 2012 Cost of antibiotic resistance and the geometry of adaptation. Mol. Biol. Evol. 29: 1417–1428.

Sprinthall, R. C., 2014 Basic statistical analysis. Pearson, 9th edition.

Tenaillon, O., 2014 The utility of fisher’s geometric model in evolutionary genetics. Annu. Rev. Ecol. Evol. Syst. 45: 179–201.

Whitehead, T. A., A. Chevalier, Y. Song, C. Dreyfus, S. J. Fleishman, C. De Mattos, C. A. Myers, H. Kamisetty, P. Blair, I. A. Wilson, and D. Baker, 2012 Optimization of affinity, specificity and function of designed influenza inhibitors using deep sequencing. Nat Biotech 30: 543–548.

Williams, D., 1991 Probability with Martingales. Cambridge mathematical textbooks, Cambridge University Press.

Wu, N. C., A. P. Young, S. Dandekar, H. Wijersuriya, L. Q. Al-Mawsawi, T.-T. Wu, and R. Sun, 2013 Systematic identification of h274y compensatory mutations in influenza a virus neuraminidase by high-throughput screening. J. Virol. 87: 1193–1199.

